# IL13 critically contributes to the antitumor activity of polyfunctional CD4^+^ T cells driven by constitutively active Stat5

**DOI:** 10.64898/2026.11.09.765236

**Authors:** Ogacheko D. Okoko, Mercy Kehinde-Ige, Xin Wang, Yan Ye, Aarav Bhasin, Lirong Pei, Caitlin Brandle, Md Y. Gazi, Zhi-chun Ding, Rafal Pacholczyk, Huidong Shi, Gang Zhou

**Author notes:** **Correspondence:** Huidong Shi, PhD Georgia Cancer Center, Augusta University, 1120 15^th^ Street, CN-2138 Augusta, GA 30912, Gang Zhou*****, PhD Georgia Cancer Center, Augusta University, 1120 15^th^ Street, CN-4140 Augusta, GA 30912.

## Abstract

The presence of polyfunctional T cells capable of concurrently producing multiple cytokines correlates with favorable clinical outcomes in patients with cancer. We previously reported that overexpression of constitutively active Stat5a (CASTAT5) confers polyfunctionality on tumor-specific CD4⁺ T cells, enabling durable antitumor immunity following adoptive cell therapy. However, the mechanisms underlying the enhanced antitumor effects of CASTAT5-modified T cells remain unclear. Here, we identify interleukin-13 (IL13) as a critical effector cytokine contributing to the multifaceted antitumor activity of CASTAT5-conditioned polyfunctional CD4⁺ T cells (C5-CD4). C5-CD4 T cells exhibit a prominent T helper 2 (Th2) cytokine profile, accompanied by extensive epigenomic remodeling at the Th2 cytokine loci, as revealed by Hi-C and histone-modification CUT&Tag assays. Neutralization of IL13 diminished the antitumor efficacy of C5-CD4 T cells. Furthermore, targeted *Il13* disruption substantially impaired tumor control and was associated with reduced donor T cell expansion and persistence, diminished systemic cytokine and chemokine responses, and attenuated activation of endogenous tumor-reactive CD8⁺ T cells. Nonetheless, IL13 overexpression alone in donor CD4⁺ T cells failed to fully recapitulate the therapeutic activity of C5-CD4 T cells, indicating that IL13 is a critical contributor but is insufficient to account for the full antitumor activity conferred by CASTAT5. Single-cell RNA sequencing associated donor-derived IL13 with increased representation of tumor-infiltrating myeloid cells and IL13-signaling in monocytes and macrophages exhibiting tissue-repair-associated transcriptional programs. Moreover, *Il13* disruption diminished the therapeutic efficacy of CASTAT5-expressing CD19-targeted CAR-T cells in a mouse B-cell lymphoma model. Reanalysis of a human CD19CAR-T infusion-product dataset further revealed an association between IL13 expression and type 2 cytokine signatures linked to prolonged clinical response, suggesting a potential contribution of IL13 to the efficacy of human CD19CAR-T therapy. Together, these findings uncover an unexpected antitumor role for IL13 and establish it as a critical node linking polyfunctional donor T cells with endogenous immune cells to promote sustained tumor immunity.

## Introduction

Adoptive T-cell therapy (ACT) has emerged as a powerful approach to cancer treatment, yet achieving durable tumor control remains challenging. Although CD8⁺ T cells have traditionally been viewed as the primary therapeutic effectors, accumulating evidence demonstrates that CD4⁺ T cells can also mediate potent and durable antitumor responses in ACT using chimeric antigen receptor-modified T cells (CAR-T) and T-cell receptor-engineered T cells (TCR-T)(*1–4*). CD4⁺ T cells can coordinate broader immune responses through cytokine production, activation of other lymphocytes, and modulation of myeloid cells, stromal cells and tumor vasculature within the tumor microenvironment (TME)(*5–12*). However, the cellular programs and effector molecules that enable CD4⁺ T cells to sustain antitumor immunity remain incompletely defined.

The presence of polyfunctional T cells capable of simultaneously producing multiple proinflammatory cytokines correlates with favorable clinical outcomes in patients with cancer(*13–16*). In particular, polyfunctional CD4⁺ T cells have been associated with enhanced antitumor immune responses in both preclinical models and cancer patients(*17–21*). We previously demonstrated that enforced expression of constitutively active Stat5a (CASTAT5) endows antigen-specific CD4⁺ T cells with a distinctive polyfunctional phenotype, enabling durable and curative antitumor activity in a murine B-cell lymphoma model(*22*). CASTAT5-conditioned polyfunctional CD4⁺ T cells (C5-CD4) simultaneously exhibit features of multiple effector programs, including Th1, Th2, and Th9 responses, as well as granulocyte-macrophage colony-stimulating factor (GM-CSF) production and cytolytic activity. Importantly, these cells persist long-term after adoptive transfer and elicit robust endogenous antitumor immunity, suggesting that their therapeutic effects extend beyond the direct activity of the transferred cells. Moreover, CASTAT5 can markedly improve the expansion and persistence of CD19CAR-T cells, resulting in a high cure rate in mice with advanced lymphoma. Nevertheless, how this distinctive polyfunctional state translates into sustained tumor immunity remains poorly understood.

The prominent Th2 component of C5-CD4 cells raises a particularly intriguing question regarding the role of type 2 immunity in antitumor responses. The primary Th2 cytokines, including IL4, IL5 and IL13, can exert context-dependent and sometimes opposing effects on tumor immunity(*23–25*). Although these cytokines have frequently been implicated in tumor-promoting inflammation, immunosuppression and tissue remodeling(*26–29*), their antitumor activities have also been well-documented(*5, 30, 31*). Findings from ACT preclinical models underscore the therapeutic potential of Th2 cells(*9, 32–34*). Recent studies have associated Th2-like transcriptional signatures in CAR-T infusion products with favorable clinical outcomes, including prolonged B-cell aplasia and durable remission(*35*). Moreover, type 2 cytokines, including IL4, IL5, and IL13, were detected at higher levels in the sera of patients with durable clinical responses. These observations suggest that type 2-associated programs may contribute to the persistence and therapeutic function of antitumor T cells. However, whether individual Th2 cytokines directly contribute to therapeutic efficacy, and how their effects depend on the broader cytokine milieu and cellular context, remain unresolved.

Here, we identify IL13 as a critical contributor to the antitumor activity of C5-CD4 polyfunctional T cells. Epigenomic analyses of C5-CD4 T cells reveal extensive chromatin remodeling at Th2 cytokine loci, providing insight into the epigenetic mechanisms underlying the prominent type 2 cytokine profile induced by CASTAT5. Using complementary genetic and pharmacological approaches, we demonstrate that IL13 is required for optimal C5-CD4-mediated tumor control but is insufficient, on its own, to recapitulate the full therapeutic activity conferred by CASTAT5. Single-cell transcriptomic analyses further associate donor T cell-derived IL13 with changes in tumor-infiltrating myeloid cell abundance, as well as with broader endogenous immune responses. We also show that *Il13* disruption attenuates the therapeutic efficacy of CASTAT5-expressing CD19CAR-T. Finally, reanalysis of a human CD19CAR-T infusion-product dataset reveals an association between IL13 expression, type 2 cytokine programs, and favorable clinical outcomes. Together, these findings uncover an unexpected antitumor role for IL13 and identify it as a key mediator linking polyfunctional donor T cells with the host immune microenvironment to support sustained antitumor immunity.

## RESULTS

### CASTAT5 induces a polyfunctional CD4⁺ T-cell state characterized by prominent Th2 cytokine production

We previously reported that overexpression of CASTAT5 endows CD4⁺ T cells with a polyfunctional phenotype and that adoptive transfer of CASTAT5-transduced, HA-specific CD4⁺ T cells induces durable, curative responses in mice bearing HA-expressing A20 B-cell lymphoma(*22*). However, those studies used bulk-transduced CD4⁺ T cells for adoptive transfer, leaving open the possibility that differences in transduction efficiency contributed to inter-experimental variability. To overcome this limitation, we generated a retroviral vector encoding CASTAT5 linked to a cistronic green fluorescent protein (GFP) reporter (CASTAT5-GFP) (Fig. 1A). This design enabled purification of CASTAT5-transduced, HA-specific CD4⁺ T cells based on GFP expression (C5-CD4), while the GFP-negative fraction from the same donor preparation served as the control population (CTRL-CD4) (Fig. 1B).

**Figure 1.**
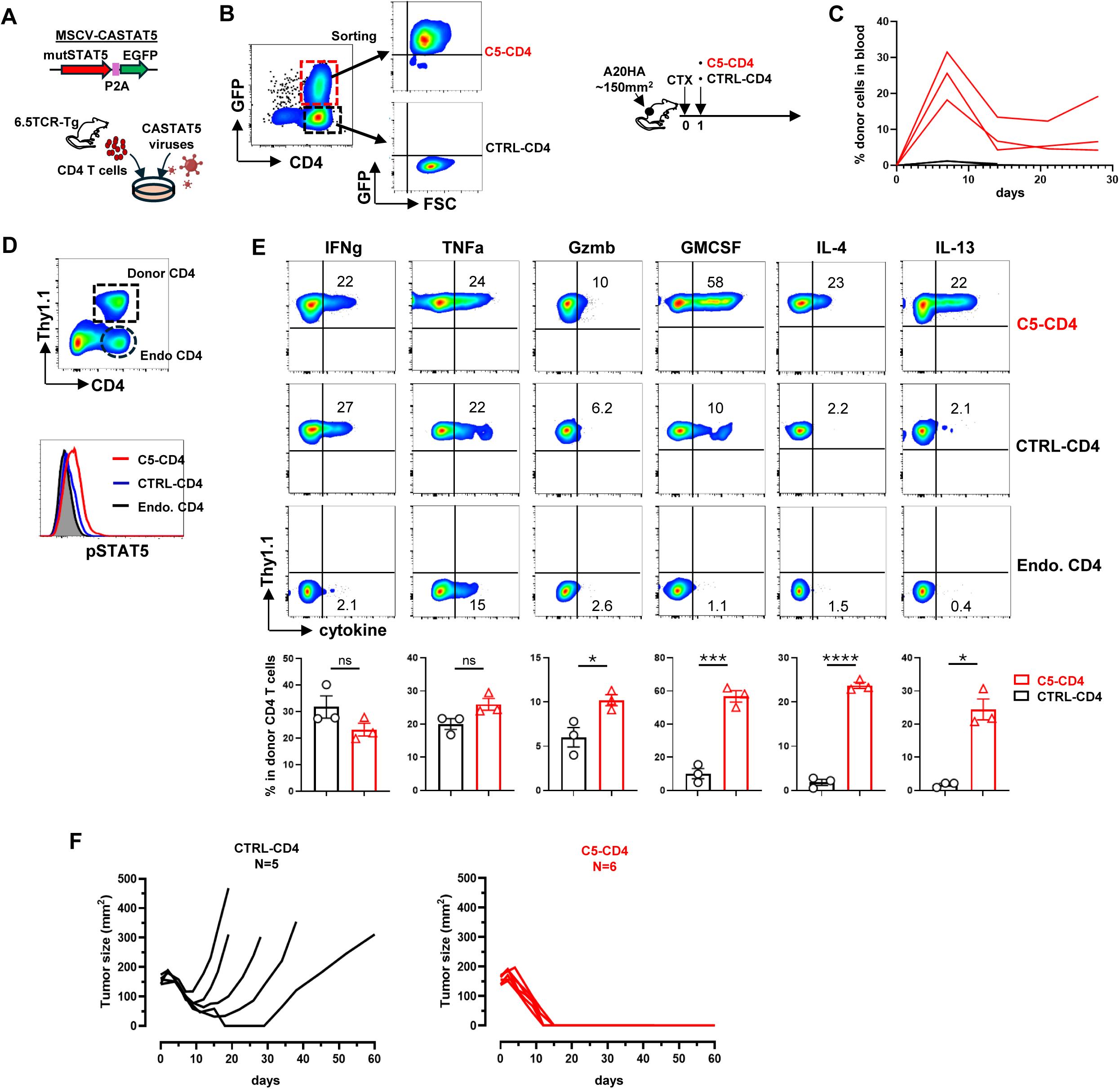
CASTAT5 induces a polyfunctional CD4 T-cell state with a prominent Th2 cytokine profile and diverse inflammatory and cytotoxic effector functions. A. Schematic of the cistronic retroviral vector encoding constitutively active STAT5 (CASTAT5) followed by a P2A-linked EGFP reporter. The vector was used to transduce CD4+ T cells isolated from splenocytes of 6.5 TCR-Tg mice. B. Schematic of the isolation and adoptive transfer of CASTAT5-transduced HA-specific CD4^+^ T cells. Transduced cells were separated by GFP expression into GFP^+^ C5-CD4 and GFP^−^ CTRL-CD4 populations by cell-sorter and adoptively transferred into mice bearing established A20HA tumors. All mice received cyclophosphamide (CTX) chemo-conditioning one day prior to T cell infusion. C. Frequencies of donor CD4^+^ T cells in peripheral blood following adoptive transfer. Tail blood samples (3 mice per group) were collected weekly, and donor cells were quantified by flow cytometry. **D.** Intracellular pStat5 expression in donor CD4^+^ T cells. Representative flow cytometry plots illustrate the gating strategy used to identify donor and endogenous CD4^+^ T cells, followed by assessment of pSTAT5 expression. **E.** Cytokine and effector profile of donor CD4^+^ T cells assessed by intracellular cytokine staining (ICS). Representative dot plots are shown for each indicated condition, with endogenous CD4^+^ T cells used to establish the gating strategy. Numbers indicate the percentages of cytokine-producing cells in donor CD4^+^ T cells. Quantification is summarized in the corresponding bar graphs shown as mean ± SE of three mice per group. **F.** Tumor growth curves of mice bearing established A20HA tumors following adoptive transfer of the indicated donor CD4^+^ T-cell populations. ns, not significant; *, p < 0.05; ***, p < 0.001; ****, p < 0.0001.

Following adoptive transfer, C5-CD4 T cells underwent robust expansion and persisted long term in recipient mice, whereas CTRL-CD4 T cells showed limited expansion and failed to persist (Fig. 1C). Consistent with constitutive Stat5 activation, C5-CD4 T cells exhibited higher pStat5 levels than CTRL-CD4 T cells (Fig. 1D). To characterize the functional phenotype of the donor cells in vivo, we performed intracellular cytokine staining (ICS) of splenic T cells 7 days after adoptive transfer, when tumor burdens remained comparable between the two treatment groups. C5-CD4 T cells displayed enhanced production of granzyme B (GzmB), GM-CSF, IL4, and IL13 compared with CTRL-CD4 T cells, whereas IFNγ and TNFα production was largely comparable between the groups (Fig. 1E). Thus, forced CASTAT5 expression generated a distinct polyfunctional CD4 T-cell phenotype characterized by enhanced cytotoxic and cytokine-producing capacity, with particularly prominent production of the Th2 cytokines IL4 and IL13.

Consistent with our previous findings, all mice receiving C5-CD4 T cells achieved complete tumor regression and remained tumor-free, whereas mice receiving CTRL-CD4 T cells experienced tumor relapse (Fig. 1F). Notably, CASTAT5-CD4 T cells remained readily detectable in the spleens of cured mice 60 days after T-cell infusion (Supplemental Fig. 1A). At this late time point, the persisting C5-CD4 T cells, rather than endogenous CD4⁺ T cells, were the predominant producers of IL4, IL13, GM-CSF, IFNγ, and TNFα and retained their polyfunctional phenotype (Supplemental Fig. 1B). Moreover, persisting C5-CD4 T cells exhibited higher levels of pStat5, BCL2, and GATA3 than endogenous CD4⁺ T cells (Supplemental Fig. 1C), consistent with sustained Stat5 signaling, enhanced survival capacity, and maintenance of a Th2-associated transcriptional program.

We next asked whether overexpression of wild-type Stat5 (WTSTAT5) could recapitulate the phenotype and antitumor activity induced by CASTAT5. To this end, equal numbers of CASTAT5-or WTSTAT5-transduced HA-specific CD4⁺ T cells were adoptively transferred into A20HA tumor-bearing mice (Supplemental Fig. 2A). ICS of donor cells in peripheral blood 7 days after T-cell infusion revealed that C5-CD4 T cells expressed significantly higher levels of pStat5, BCL2, GMCSF, IL4 and IL13 than WTSTAT5-CD4 T cells, whereas IFNγ and TNFα production did not differ significantly between the two donor cell populations (Supplemental Fig. 2B). In addition, WTSTAT5-CD4 T cells were less effective than C5-CD4 T cells in controlling tumor growth and failed to achieve comparable rates of complete tumor regression (Supplemental Fig. 2C).

### CASTAT5 drives chromatin remodeling at Th2 cytokine loci

The ability of donor CASTAT5-CD4 T cells to persist and retain polyfunctionality long after tumor eradication, as shown in Supplemental Fig. 1, suggests that CASTAT5 may establish durable epigenetic programs that sustain their functional phenotype. Our previous analysis revealed significant changes in chromatin accessibility across multiple T-cell programs, including regulatory regions associated with cytokine genes(*22*). To further characterize CASTAT5-mediated epigenomic remodeling in polyfunctional tumor-reactive CD4⁺ T cells, we performed Hi-C and H3K27ac and H3K27me3 CUT&Tag sequencing on donor C5-CD4 T cells recovered from long- term surviving mice (60 days post T cell transfer), using endogenous CD4⁺ T cells from the same mice as controls. We integrated the resulting Hi-C contact maps and histone-modification profiles with a published Stat5-binding landscape in activated CD4⁺ T cells to investigate the relationship between CASTAT5-associated chromatin remodeling and Stat5 occupancy (Fig. 2A).

**Figure 2.**
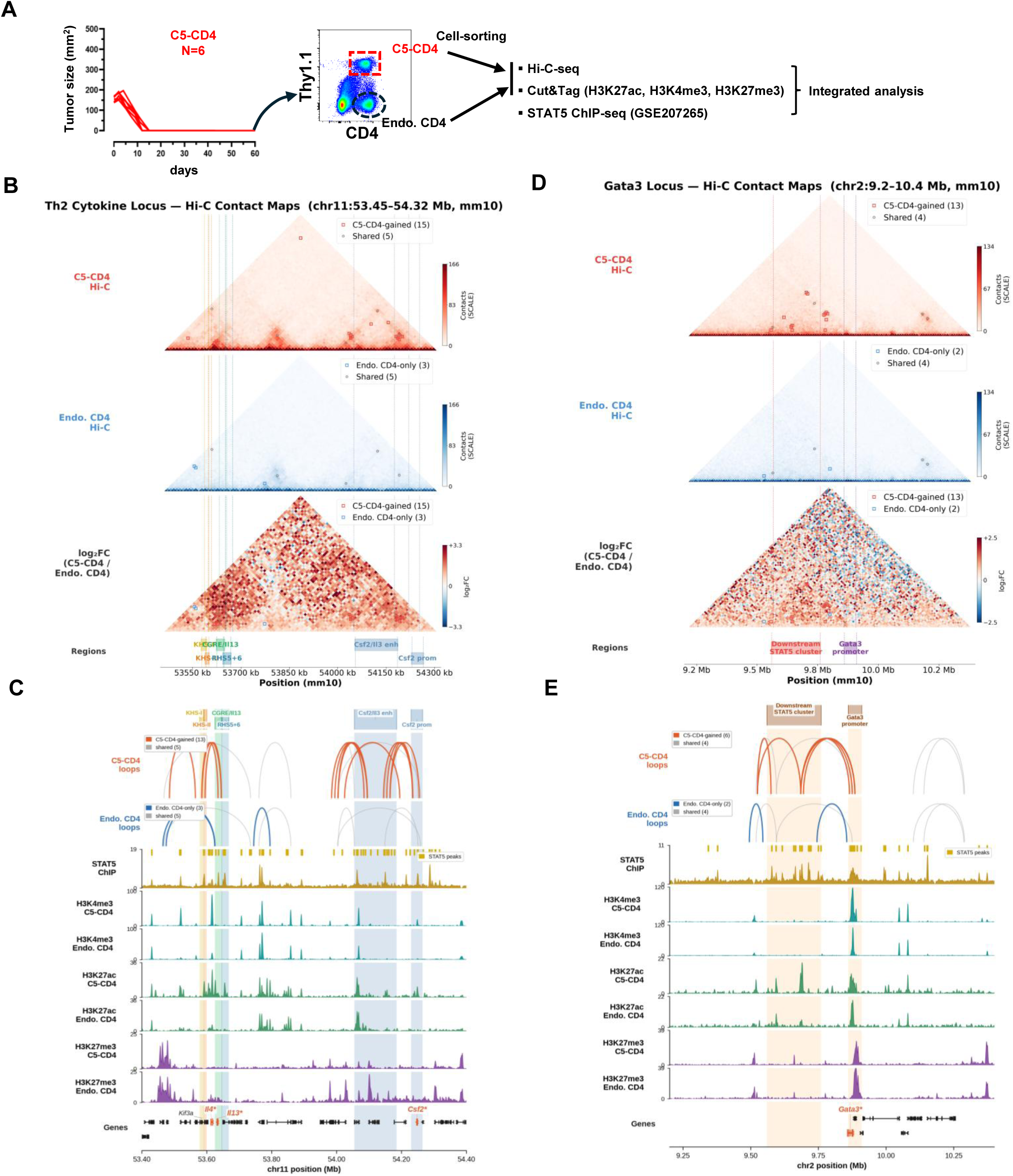
CASTAT5 is associated with changes in chromatin contacts and histone modifications at the Th2 cytokine locus and the Gata3 locus. A. Experimental design. Donor C5-CD4 T cells and endogenous CD4^+^ T cells (Endo. CD4) were sorted from the spleens of mice 60 days following C5-CD4 transfer and profiled by Hi-C and CUT&Tag (H3K4me3, H3K27ac, and H3K27me3). Reference Stat5 ChIP- seq data are from IL-2-stimulated mouse CD4^+^ T cells (GSE207265). B. Hi-C contact maps of the Th2 cytokine locus (chr11:53.45–54.32 Mb, mm10) at 10-kb resolution, SCALE-normalized with Juicer Tools, for C5-CD4 and Endo. CD4 T cells, with the log₂ contact ratio log₂[(C5-CD4 + 1)/(Endo. CD4 + 1)]. Color scales are capped at the 99th percentile of contacts within 500 kb of the diagonal. Squares mark condition-specific HiCCUPS loop calls, and circles mark shared contacts. Counts in the keys are full-depth calls with both anchors inside the displayed window; a call was classified as shared if the other condition had a call within ±1 bin at both anchors, and shared calls from the two conditions were merged into one contact. Shaded intervals mark the Kif3a-associated hypersensitive sites KHS-I and KHS-II, the conserved Gata3 response element (CGRE) upstream of Il13, hypersensitive sites RHS5–RHS6 of the Th2 locus control region, the Csf2/Il3 regulatory interval and the Csf2 promoter. C. Genome-browser view of chr11:53.4–54.4 Mb showing loop arcs, reference Stat5 ChIP-seq signal and peaks, and H3K4me3, H3K27ac, and H3K27me3 CUT&Tag signal in C5-CD4 and Endo. CD4 T cells. Only loops that remained condition-specific after C5-CD4 contacts were down-sampled to Endo. CD4 depth and loops were re-called with HiCCUPS are drawn: 13 of 15 C5-CD4- gained loops, 3 of 3 Endo. CD4-only loops and 5 of 5 shared contacts. D. Hi-C contact maps of the Gata3 locus (chr2:9.2–10.4 Mb), as in B. Shaded intervals mark a downstream cluster of reference Stat5 peaks and the Gata3 promoter region. E. Genome-browser view of the Gata3 locus, as in C; 6 of 13 C5-CD4-gained loops, 2 of 2 Endo. CD4-only loops and 4 of 4 shared contacts remained at matched depth. In C and E, CUT&Tag tracks show the mean of three biological replicates per mark and condition, and the Stat5 track shows the mean of two ChIP-seq replicates; signal was averaged into 1,600 bins across each window, and each mark is drawn on a common y-axis for both conditions, capped at the 99.5th percentile of the pooled signal.

Our previous ATAC-seq analysis identified the *Il4–Il13* locus as one of the regions showing the most pronounced changes in chromatin accessibility in C5-CD4 donor cells(*22*). Hi-C analysis further revealed extensive gains in chromatin contacts across the extended Th2 cytokine region (approximately 1 Mb in length, chr11:53.4–54.4 Mb) in C5-CD4 donor cells relative to endogenous CD4⁺ T cells (Fig. 2B). This region encompasses several well-characterized cis-regulatory elements, including the conserved GATA3 response element (CGRE) near *Il13*(*36*), RHS5 and RHS6 within the Th2 locus control region (LCR)(*37*), the *Kif3a*-associated hypersensitive sites KHS-I/II implicated in chromatin organization(*38*), and upstream regulatory elements associated with the *Csf2/Il3* locus(*39*). At full sequencing depth, Hi-C identified 15 loops specific to C5-CD4 cells and three specific to endogenous CD4⁺ T cells (Fig. 2B). Because the C5-CD4 library was sequenced more deeply, we down-sampled C5-CD4 contacts to the depth of the endogenous

CD4 library and re-called loops. Thirteen of the 15 C5-CD4-specific loops and all three endogenous-specific loops were retained (Fig. 2C). Three of the gained loops connected the *Kif3a*-embedded KHS-I/II region to the *Il13* promoter–CGRE region, and four connected a regulatory interval located approximately 60–190 kb 3′ of *Csf2* to the *Csf2* locus. Integration with the published STAT5-binding dataset(*40*) revealed STAT5 occupancy within 5 kb of both loop anchors for 9 of the 13 gained loops. H3K27ac profiling identified 13 gained peaks across the region, 10 of them assigned to *Kif3a*, *Il4*, *Il13*, or *Csf2* (Fig. 2C). Six of the retained loops showed increased H3K27ac at one or more anchors, including four with increased H3K27ac at both anchors, among them all three KHS–Il13 loops. H3K4me3 was also increased at the Il4 promoter (log2FC = 3.31, adjusted P = 3 × 10⁻¹⁸) and, nominally, at the *Csf2* promoter (log2FC = 3.44, P = 0.005). Together, these findings reveal coordinated changes in active histone marks and chromatin contacts at regulatory elements with reference STAT5 occupancy across the Th2 cytokine region. These changes are consistent with the increased production of IL4, IL13, and GM-CSF by C5-CD4 T cells measured by flow cytometry (Fig. 1E) and suggest that CASTAT5-associated chromatin remodeling contributes to the enhanced expression of these cytokines.

Given its key role in Th2 differentiation, we next examined the *Gata3* locus (chr2:9.2-10.4 Mb). At full sequencing depth, Hi-C identified 13 loops specific to C5-CD4 cells, 2 loops specific to endogenous CD4⁺ cells, and 4 shared contacts (Fig. 2D). After down-sampling the C5-CD4 contacts to the depth of the endogenous CD4 library, 6 of the 13 C5-CD4-specific loops and both endogenous-specific loops were retained (Fig. 2E). Each of the retained C5-CD4-specific loops had one anchor within a cluster of 15 reference STAT5 occupancy peaks located ∼120-320 kb downstream of the *Gata3* TSS (chr2:9.56-9.76 Mb). Four of the loops connected the cluster to the *Gata3* promoter, and three connected to an anchor that contained the *Gata3* TSS. The remaining two connected a site at chr2:9.52 Mb to the cluster. All six loops had reference STAT5 occupancy within 5 kb of at least one anchor, and for the four cluster-to-promoter loops, occupancy was present at both anchors. Five of the six loops showed increased H3K27ac at one or more anchors (including the three loops connecting the cluster to the *Gata3* TSS, which showed increased H3K27ac at both anchors). Neither of the endogenous-specific loops reached the *Gata3* promoter. At the *Gata3* promoter, C5-CD4 cells showed increased H3K4me3 (log2FC = 2.25, adj *P* = 1.6 x 10⁻^8^) and decreased repressive H3K27me3 (log2FC = -1.01, adj *P* = 0.028). H3K27ac was also higher (log2FC = 1.83, *P* = 0.002), though this did not survive FDR correction (adj *P* = 0.08). The gains in H3K4me3 were restricted to the promoter-side anchors. In summary, these data associate CASTAT5 with depth-robust contacts between a downstream STAT5-occupied region and the *Gata3* promoter, with a more active histone profile at the promoter consistent with the higher levels of GATA3 protein observed in the persistent C5-CD4 cells (Suppl Fig. 1C).

### IL13-deficiency impairs the therapeutic activity of C5-CD4 T cells

The polyfunctional phenotype of C5-CD4 T cells, characterized by the simultaneous production of cytokines associated with distinct T-cell effector programs, prompted us to investigate the functional contributions of cytokines prominently produced by these cells. Given the pronounced Th2-associated transcriptional and epigenetic profiles of these cells, we examined whether neutralizing IL4 or IL13 affected their therapeutic activity. Administration of an IL13-neutralizing antibody following C5-CD4 T-cell transfer impaired antitumor efficacy, resulting in increased tumor relapse and reduced long-term survival; in contrast, neutralization of IL4 did not compromise therapeutic efficacy (Supplemental Fig. 3A-B). The results implicate IL13 as a key Th2 cytokine contributing to the antitumor activity of C5-CD4 T cells.

To further determine the role of donor-derived IL13, we used CRISPR-Cas9 genome editing to disrupt *Il13* in C5-CD4 T cells. Seki et al. previously described an optimized Cas9 ribonucleoprotein (RNP)-based protocol that enables highly efficient genome editing in primary mouse T cells, with near-complete loss of target gene expression at the population level(*41*). We adapted this approach using the Alt-R CRISPR-Cas9 genome-editing system (IDT), which employs synthetic single-guide RNAs (sgRNAs) complexed with recombinant Cas9 protein, to disrupt *IL13* in C5-CD4 T cells. Before targeting *IL13*, we first established the efficiency and compatibility of this approach with our C5-CD4 T-cell generation protocol by targeting CD90.1 (Thy1.1), a congenic surface marker expressed by CD4^+^ T cells from HA-TCR Tg mice. As outlined in Supplemental Fig. 4A, nucleofection of activated CTRL-CD4 and C5-CD4 T cells with sg*CD90.1*/Cas9 RNP resulted in greater than 90% loss of CD90.1 expression in both populations (Supplemental Fig. 4B). This result established an efficient RNP-based editing strategy compatible with C5-CD4 T-cell generation. We subsequently targeted *Il13* in C5-CD4 T cells. Nucleofection with sg*Il13*/Cas9 RNP did not affect CASTAT5 transduction efficiency, as assessed by GFP expression (Supplemental Fig. 4C), or intracellular pStat5 levels (Supplemental Fig. 4D). Editing of the *Il13* locus was demonstrated by T7 endonuclease I digestion (Supplemental Fig. 4E) and independently confirmed by nanopore sequencing (Supplemental Fig. 4F).

To assess the functional consequences of IL13 deficiency in C5-CD4 T cells, equal numbers of sg*Il13*C5-CD4 and C5-CD4 T cells were adoptively transferred into A20HA-bearing mice. Donor T-cell frequencies and cytokine-producing capacity, endogenous CD8 T-cell activation, circulating cytokine profiles, and antitumor efficacy were assessed at the indicated time points to determine how IL13 loss affected the immune and therapeutic functions of C5-CD4 T cells (Fig. 3A). sg*Il13*C5-CD4 T cells were readily detectable at day 7, with frequencies ranging from 9% to 45%, but declined to barely detectable levels by day 21; in contrast, the abundance of C5-CD4 T cells remained relatively stable over this period (Fig. 3B). Notably, cytokine ICS of donor T cells in blood collected on day 7 revealed that sg*Il13*C5-CD4 T cells exhibited a marked reduction in IL13 production, with only ∼10% of donor cells producing IL13 compared with ∼55% in C5-CD4 T cells; In contrast, production of other cytokines was either modestly affected (IFNγ and IL4) or remained unchanged (TNFα and GM-CSF) between the two donor cell populations (Fig. 3C). Because T cells were stimulated in vitro with PMA and ionomycin for 4 hours before ICS, these measurements reflect their cytokine-producing capacity under strong, nonphysiological stimulation rather than their cytokine production in vivo. These results confirm effective functional disruption of IL13 while largely preserving the capacity of sg*Il13*C5-CD4 T cells to produce other cytokines.

**Figure 3.**
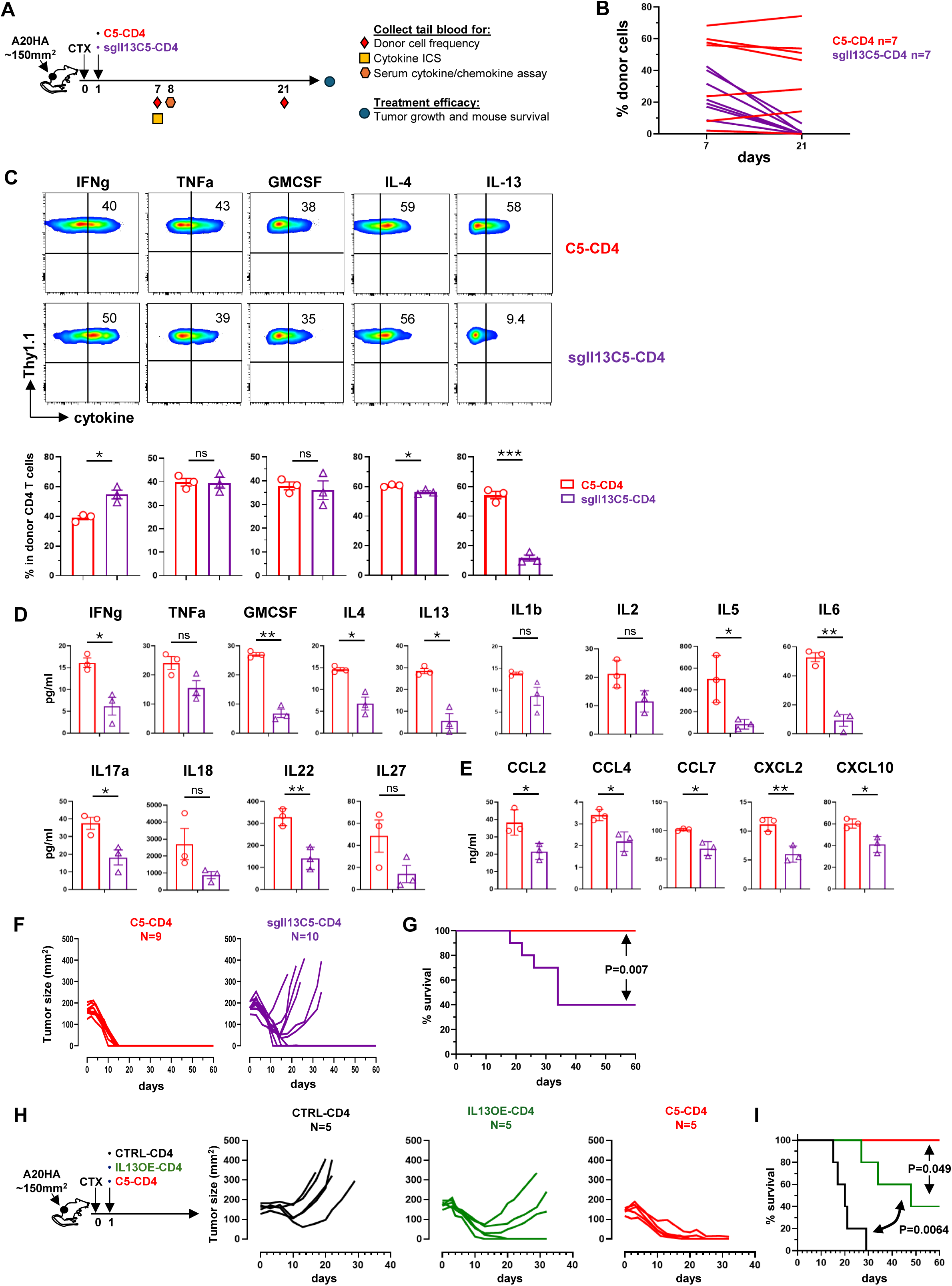
IL13-deficiency in C5-CD4 T cells alters donor cell expansion and phenotype, the systemic inflammatory milieu, and antitumor efficacy. A. Schematic of the experimental timeline. Following adoptive transfer of C5-CD4 and sgIl13C5-CD4 T cells, peripheral blood was collected by tail vein at the indicated time points for the specified analyses. Mice were monitored for tumor growth and survival. B. Frequencies of donor T cells in peripheral blood on days 7 and 21 after adoptive transfer, determined by flow cytometry. C. Cytokine profile of donor CD4+ T cells assessed by ICS. Representative dot plots are shown for each condition, with numbers indicating the percentages of cytokine-producing cells among donor CD4+ T cells. Quantification is summarized in the corresponding bar graphs as mean ± SE (n = 3 mice per group). D-E. Systemic cytokine and chemokine milieu assessed by Luminex on day 8. Concentrations of the indicated cytokines and chemokines are shown as mean ± SE (n = 3 mice per group). F. Tumor growth curves for individual mice in each treatment group. G. Kaplan–Meier survival analysis of mice shown in F. H. Comparison of the antitumor efficacy of adoptively transferred IL13-overexpressing CD4 T cells (IL13OE-CD4), C5-CD4 T cells, and CTRL-CD4 T cells. The experimental groups are depicted schematically. Tumor growth curves for individual mice are shown. I. Kaplan–Meier survival analysis of mice shown in H. ns, not significant; *, p < 0.05; **, p < 0.01; ***, p < 0.001.

Despite the relatively preserved cytokine-producing capacity observed by ICS, selective loss of IL13 in donor T cells was associated with broader changes in the systemic cytokine milieu. Measurement of serum cytokines, which provides a snapshot of the circulating inflammatory environment *in vivo*, revealed significantly reduced levels of IFNγ, GM-CSF, IL4, IL13, IL5, IL6, IL17A, and IL22 in mice receiving sg*Il13*C5-CD4 T cells compared with those receiving control C5-CD4 T cells (Fig. 3D). TNFα, IL1β, IL2, IL18, and IL27 levels were also reduced, although these differences did not reach statistical significance. In addition, mice receiving sg*Il13*C5-CD4 T cells exhibited reduced circulating levels of inflammatory chemokines, including CCL2, CCL4, CCL7, CXCL2, and CXCL10 (Fig. 3E). Thus, loss of IL13 in C5-CD4 donor T cells was associated with a pronounced attenuation of the systemic inflammatory cytokine and chemokine milieu.

At the therapeutic endpoint, sg*Il13*C5-CD4 T cells exhibited substantially reduced antitumor efficacy compared with C5-CD4 T cells. Six of 10 mice receiving sg*Il13*C5-CD4 T cells experienced tumor relapse followed by early death, whereas all mice receiving C5-CD4 T cells (9 of 9) achieved complete tumor regression and prolonged survival (Fig. 3F-G). Because sg*Il13*-mediated editing could potentially impair the overall fitness or functional state of C5-CD4 T cells independent of IL13 loss, we next tested whether the nucleofection and CRISPR manipulation procedures themselves affected therapeutic activity. To this end, we generated C5-CD4 T cells nucleofected with a scrambled sgRNA (sgScrC5-CD4) and compared their antitumor activity with that of non-nucleofected C5-CD4 T cells (Supplemental Fig. 5A). Nucleofection with scramble sgRNA did not affect CASTAT5 transduction efficiency (Supplemental Fig. 5B), and adoptive transfer of sgScrC5-CD4 T cells was equally effective as the transfer of non-nucleofected C5-CD4 T cells in achieving complete tumor regression (Supplemental Fig. 5C). These controls indicate that the impaired therapeutic activity following *Il13* targeting was not attributable to the editing procedure itself.

We next asked whether IL13 overexpression alone could recapitulate the antitumor activity conferred by CASTAT5. To this end, HA-specific CD4⁺ T cells were transduced with a retroviral vector encoding murine IL13 and the mCherry reporter (Supplemental Fig. 6A-B). The resulting IL13-overexpressing CD4⁺ T cells (IL13OE-CD4), mock-transduced CD4⁺ T cells (CTRL-CD4), and C5-CD4 T cells were then compared in adoptive transfer experiments (Fig. 3H schema). Although IL13OE-CD4 T cells inhibited tumor growth more effectively than CTRL-CD4 T cells, their therapeutic efficacy was inferior to that of C5-CD4 T cells, particularly in terms of long-term survival (Fig. 3H-I). Moreover, IL13OE-CD4 donor cells were no longer detectable in peripheral blood three weeks after transfer, whereas C5-CD4 donor cells remained readily detectable (Supplemental Fig. 6C), suggesting that IL13 overexpression alone was insufficient to sustain donor cell persistence.

Collectively, these findings demonstrate that IL13 is a critical contributor to the antitumor activity of C5-CD4 T cells but is insufficient on its own to recapitulate the full therapeutic effects conferred by CASTAT5.

### scRNA-seq analysis reveals transcriptomic changes in IL13-deficient C5-CD4 T cells

To investigate how donor-derived IL13 influences the tumor immune microenvironment (TME), we performed single-cell RNA sequencing (scRNA-seq) of tumor-infiltrating CD45+ immune cells from mice receiving C5-CD4, mock-transduced CTRL-CD4, or sg*Il13*C5-CD4 T cells (Fig. 4A). In total, 45,097 cells were profiled across the three groups (Fig. 4B), encompassing T cells, myeloid cells, other immune populations, and tumor cells (B-cell lymphoma). Each condition was represented by a single pooled sequencing sample; therefore, comparisons describe differences among the recovered cell populations and should be interpreted as descriptive rather than as biological replicate-level statistical comparisons. Compared with the other two groups, tumors from C5-CD4-treated mice contained a lower proportion of T cells but higher proportions of NK cells and myeloid cells (Fig. 4C).

**Figure 4.**
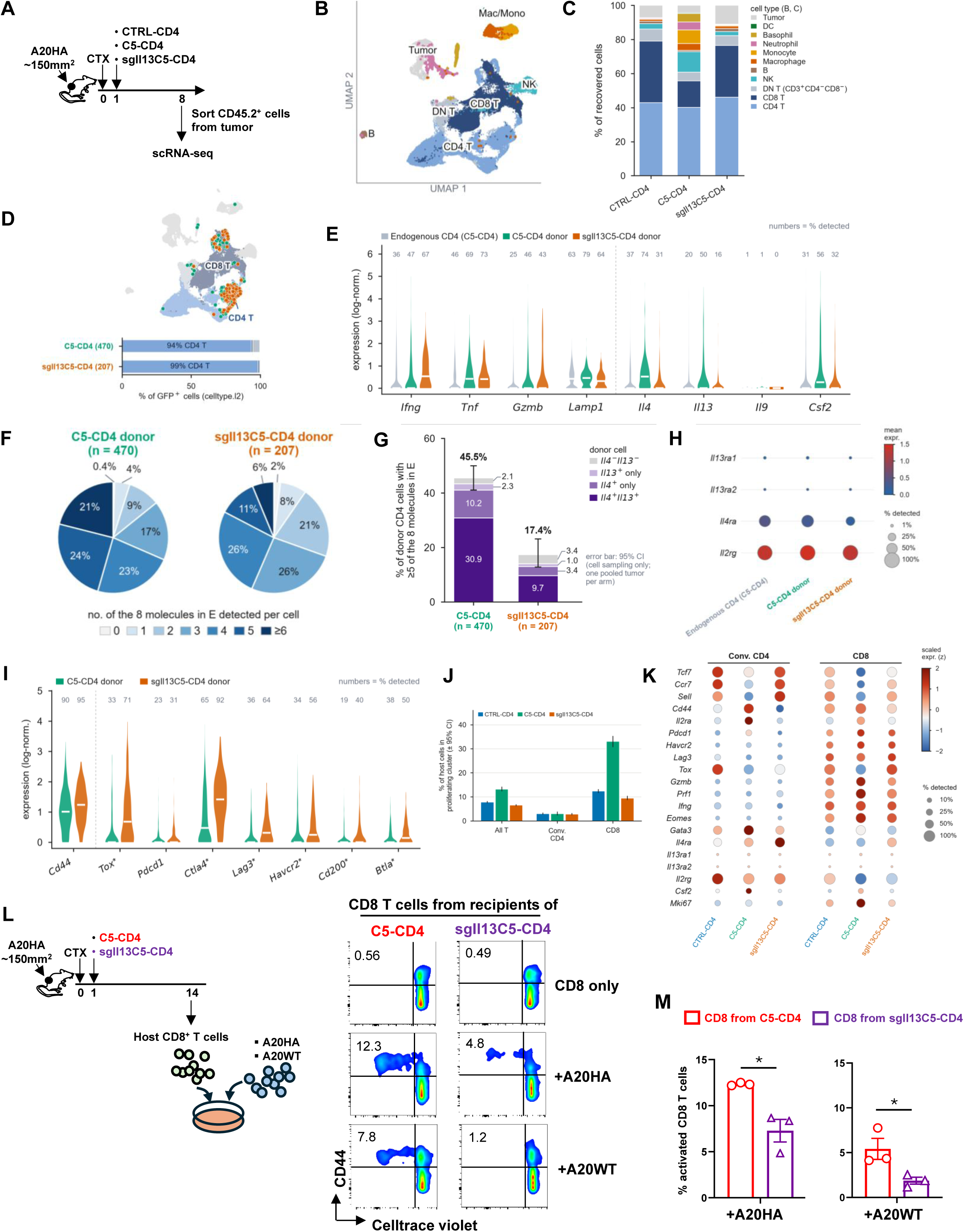
IL13 deficiency alters C5-CD4 T-cell polyfunctionality and is associated with phenotypical changes in endogenous T cells. **A.** Schematic of the experimental design for scRNA-seq analysis of immune cells isolated from A20HA tumors following adoptive transfer of control CD4^+^ T cells (CTRL-CD4), CASTAT5-CD4 T cells (C5-CD4), or CASTAT5-CD4 T cells with Il13 targeted by sgRNA (sg*Il13*C5-CD4). **B.** UMAP visualization of tumor-infiltrating cells annotated by cell type. **C.** Relative abundance of major cell populations across treatment groups. **D.** UMAP highlighting GFP⁺ donor CD4 T cells (C5-CD4 or sgIl13C5-CD4) and their annotated cell identities. **E.** Expression of cytokines and effector genes (*Ifng*, *Tnf*, *Gzmb*, *Lamp1, Il4*, *Il13*, *Il9*, *Csf2*) in endogenous and donor CD4^+^ T cells. Violin plots show log-normalized gene expression, with white bars indicating medians and percentages indicating the proportions of cells with detectable transcripts. **F.** Pie chart showing changes in polyfunctionality of donor CD4^+^ T cells, assessed by co-expression of the eight genes shown in E. Gene positivity was determined from raw sequencing reads. The number of cells with positive expression of each effector molecule was counted based on the presence or absence of raw sequencing reads mapped to the mRNA sequences of each relevant gene. Cells with at least one read mapped to the relevant gene were denoted positive for the corresponding gene. **G.** Proportion of donor CD4^+^ T cells co-expressing at least five of the eight effector and cytokine genes shown in E, stratified by *Il4* and *Il13* expression status (double-positive, single-positive or double-negative). Stacked error bars are Wilson score 95% confidence intervals for the proportion of donor cells co-expressing ≥5 of the eight molecules; they reflect sampling of cells within each experimental condition. **H.** Dotplots showing expression of IL13 and IL4 receptor genes (*Il4ra*, *Il13ra1*, *Il13ra2, Il2rg*) in the donor T cells**. I.** Expression of inhibitory receptor and T-cell dysfunction-associated genes in donor CD4^+^ T cells. Asterisks denote genes meeting the indicated cell-level differential-expression criteria (Benjamini–Hochberg-adjusted P < 0.05, Δ ≥ 10 percentage points and |log₂ fold change| ≥ 0.5). **J.** Proliferative activity of endogenous CD4^+^ and CD8^+^ T cells, assessed by proliferating-cluster membership. Bars indicate proportions with cell-bootstrap 95% confidence intervals. **K.** Dot plot showing the expression of functional genes in endogenous CD4 T cells and CD8 T cells. Dot size represents the percentage of cells expressing each gene, and color indicates scaled mean expression (z-score). **L.** IL13-deficiency in C5-CD4 T cells diminishes the activation of endogenous CD8^+^ T cells. Following the timeline depicted in the schema, purified, violet dye-labeled host CD8^+^ T cells were incubated with an equal number of irradiated A20HA or A20WT cells. 7 days later, CD8^+^ T cells were evaluated for cell division (violet dye dilution) and activation (CD44) by flow cytometry. Representative dot plots are shown. The numbers indicate the percentage of cells in the corresponding quadrant. **M.** The frequencies of divided CD8^+^ T cells under each culture condition shown in L are summarized in bar graphs and shown as mean ± SE of triplicate cultures. Data shown are representative of two independent experiments.

We first examined whether IL13 disruption altered the transcriptional profiles of donor T cells. Based on GFP expression in the annotated single-cell dataset, we identified 470 C5-CD4 and 207 sg*Il13*C5-CD4 donor T cells (Fig. 4D), most of which were annotated as CD4^+^ T cells. Compared with C5-CD4 donor cells, sg*Il13*C5-CD4 cells showed a lower frequency of IL13- expressing cells (approximately 50% versus 16%), accompanied by reduced frequencies of *Il4*- expressing cells (74% versus 31%) and *Csf2*-expressing cells (56% versus 32%) (Fig. 4E). Within the donor-cell populations, *Il13* disruption was associated with reduced expression of *Il13* (10-fold), *Il4* (2.7-fold), and *Csf2* (4.5-fold), while *Ifng, Tnf, Gzmb,* and *Lamp1* expression remained largely unchanged. These findings indicate that IL13 disruption preferentially affected the Th2-associated and GM-CSF transcriptional program rather than broadly suppressing effector-gene expression.

To assess donor-cell polyfunctionality at the single-cell level, we quantified the number of effector-cytokine transcripts co-expressed by individual donor CD4⁺ T cells across the eight genes shown in Fig. 4E. C5-CD4 donor cells exhibited a highly polyfunctional transcriptional profile, with 45.5% of cells expressing at least five transcripts and 21.1% expressing at least six transcripts (Fig. 4F). In contrast, sg*Il13*C5-CD4 donor cells showed reduced polyfunctionality, with 17.4% of cells expressing at least five transcripts and 6.3% expressing at least six transcripts. To determine whether this reduction reflected changes in the composition of polyfunctional donor cell subsets, we further classified cells expressing at least five effector transcripts according to their *Il4* and *Il13* expression status (Fig. 4G). Among C5-CD4 donor cells, Il4^+^Il13^+^ cells accounted for 30.9% of all donor cells, while Il4^+^-only polyfunctional cells accounted for 10.2%. Following *Il13* disruption, these populations decreased to 9.7% and 3.4%, respectively. Notably, among the residual Il4^+^Il13^+^ cells, 87% of sg*Il13*C5-CD4 cells expressed at least five effector transcripts, compared with 70% of C5-CD4 cells. Thus, the overall reduction in donor-cell polyfunctionality was associated primarily with a decreased abundance of Th2-associated polyfunctional subsets, rather than a generalized loss of effector-transcript co-expression among the remaining Il4^+^Il13^+^ cells. We also examined the expression levels of IL4 and IL13 receptor genes in the donor cells and found that IL13 receptors (Il13ra1 and Il13ra2) are not expressed in donor CD4⁺ T cells, and that loss of Il13 did not change the receptor status (Fig. 4H).

Our previous study showed that CASTAT5 reduced the expression of multiple immune checkpoint molecules and exhaustion-associated transcription factors in C5-CD4 donor(*22*). In the current dataset, *Il13* disruption was associated with increased expression of several inhibitory- receptor and exhaustion-associated transcripts. Compared with C5-CD4 donor cells, sg*Il13*C5-CD4 T cells expressed higher levels of *Tox* (4.0-fold), *Cd200* (2.5-fold), *Ctla4* (2.4-fold), and *Lag3* (2.2-fold), with more modest increases in *Btla* (1.7-fold) and *Havcr2* (1.5-fold). *Pdcd1* expression was largely unchanged (1.4-fold), as was expression of the activation marker *Cd44* (1.2-fold) (Fig. 4I). These findings suggest that IL13 disruption is associated with increased expression of inhibitory-receptor and exhaustion-associated genes in recovered donor cells, despite the retention of an activation-associated transcriptional marker.

### IL13-deficiency in donor C5-CD4 T cells attenuates endogenous T-cell and NK-cell responses and is associated with reduced tumor-cell metabolic stress

We next examined endogenous CD4^+^ and CD8^+^ T cell populations across the three treatment groups. Among endogenous CD8^+^ T cells, the proportion of cycling cells was markedly higher in the C5-CD4 group (33%) than in the CTRL-CD4 (12%) or sg*Il13*C5-CD4 (9%) groups (Fig. 4J). Consistent with this proliferative phenotype, endogenous CD8 T cells from the C5-CD4 group exhibited the highest expression of the cytotoxic effector genes *Gzmb* and *Prf1*, the transcription factor *Eomes*, and the proliferation marker *Mki67*, together with reduced expression of naïve/memory-associated genes, including *Tcf7, Ccr7,* and *Sell* (Fig. 4K). A similar pattern was observed in endogenous conventional CD4^+^ T cells, which showed increased expression of *Il2ra, Cd44, Gata3,* and *Csf2* and reduced expression of *Ccr7, Sell,* and *Tcf7*. These transcriptional patterns were largely attenuated following *Il13* disruption in donor cells. Together, these findings suggest that donor-derived IL13 contributes to the activation and proliferative responses of endogenous T cells associated with C5-CD4 adoptive cell therapy.

Our previous study demonstrated that endogenous CD8^+^ T cells become primed against tumor-associated antigens and contribute to tumor rejection following adoptive transfer of C5-CD4 T cells(*22*). We therefore investigated whether donor-derived IL13 contributes to the induction of tumor-reactive endogenous CD8 T cells. Endogenous CD8 T cells were isolated from mice 14 days after adoptive transfer of C5-CD4 or sg*Il13*C5-CD4 T cells, when tumors in the C5- CD4 group had undergone nearly complete regression, whereas tumors in the sg*Il13*C5-CD4 group were beginning to relapse (Fig. 3F). Purified endogenous CD8^+^ T cells were labeled with a violet proliferation dye and co-cultured with A20HA cells or parental A20WT cells, which lack HA expression (Fig. 4L schema). Tumor-reactive CD8^+^ T-cell responses were assessed by dye dilution as a measure of proliferation and by CD44 upregulation. After seven days of co-culture, the frequency of responding CD8^+^ T cells was significantly higher in cultures using cells from C5-CD4-treated mice than in those using cells from sg*Il13*C5-CD4-treated mice when stimulated with A20HA cells (12.3 ± 0.2% versus 7.3 ± 2.1%, respectively; Fig. 4L-M). A similar difference was observed in responses to A20WT cells: 5.4 ± 2.0% of endogenous CD8 T cells from C5-CD4-treated mice responded to A20WT cells, whereas the corresponding response in the sg*Il13*C5- CD4 group was only 1.8 ± 0.7%. The response to A20WT cells suggests that endogenous CD8^+^ T cells were primed against additional A20-associated antigens beyond HA, consistent with epitope spreading. Thus, these functional assays support a role for donor-derived IL13 in promoting endogenous CD8^+^ T cell responses against both the targeted HA antigen and potentially additional tumor-associated antigens.

NK cells also constituted a greater proportion of recovered cells in the C5-CD4 group (11.9%) than in the CTRL-CD4 (3.3%) or sg*Il13*C5-CD4 (2.3%) groups (Fig. 4C). NK cells from the C5-CD4 group also exhibited higher expression of the cytotoxic effector genes *Gzmb* and *Prf1* than those from either comparator group (Supplemental Fig. 7). Median log-normalized *Gzmb* expression was 2.35 in the C5-CD4 group, compared with 1.03 and 1.12 in the CTRL-CD4 and sg*Il13*C5-CD4 groups, respectively. Similarly, median *Prf1* expression was 2.14 in the C5-CD4 group, compared with 1.24 and 1.33 in the corresponding groups. Collectively, these findings indicate that C5-CD4 adoptive cell therapy is associated with increased activation and proliferation of endogenous T cells, enhanced representation of NK cells, and elevated expression of cytotoxic effector genes. The attenuation of these responses following *Il13* disruption further supports a role for donor-derived IL13 in coordinating broader antitumor immune responses.

We next examined whether *Il13* disruption in donor C5-CD4 T cells was associated with changes in the tumor-cell state. We reanalyzed A20 lymphoma cells in the scRNA-seq dataset, comprising 3,680 cells across the three treatment groups. InferCNV copy number analysis revealed chromosomal copy number changes in the tumor-cell population, supporting their malignant identity (Supplemental Fig. 8). Tumor cells accounted for 4.6% of captured cells in C5-CD4-treated mice, compared with 7.6% in CTRL-CD4-treated mice and 11.2% in sg*Il13*C5-CD4-treated mice (Supplemental Fig. 9A). These percentages represent the fraction of captured cells rather than direct measurements of absolute tumor burden. The remaining tumor cells in the C5- CD4 group exhibited a pronounced glycolytic and hypoxia-associated stress signature, characterized by increased expression of *Gapdh, Ldha, Pkm, Pgk1, Eno1, Bnip3l, Adm,* and *Atf3*. The corresponding module score was 0.50, compared with 0.08 in CTRL-CD4 tumors (Supplemental Fig. 9B-E). This pattern was accompanied by reduced expression of mature B-cell identity genes, including *Pax5, Ebf1, Cd79a, Cd79b, Ms4a1,* and *Bach2*, with module scores of 0.19 and 0.72, respectively (Supplemental Fig. 9D and E). Both transcriptional patterns were largely reversed in tumors from sg*Il13*C5-CD4-treated mice. Tumor cells from this group exhibited a lower stress-associated module score (0.22) and a higher B-cell identity score (0.77). Moreover, A20 lymphoma cells expressed little or no *Il13ra1* or *Il13ra2*, arguing against direct IL13 signaling in the tumor cells (Supplemental Fig. 9F). Together, these findings associate C5-CD4 therapy with a tumor-cell state characterized by increased metabolic stress, reduced expression of mature B-cell identity genes, and a lower proliferative fraction. The reversal of these features following donor *Il13* disruption, together with the low expression of IL13 receptor transcripts in A20 cells, is consistent with an indirect contribution of donor-derived IL13 through changes in the surrounding tumor immune microenvironment.

### IL13 deficiency in donor C5-CD4 T cells attenuates myeloid-cell accumulation and tissue-repair-associated programs in tumors

One prominent finding from our scRNA-seq analysis was the increased representation of myeloid cells in tumors from C5-CD4-treated mice compared with CTRL-CD4-treated mice, an effect that was markedly attenuated following *Il13* disruption in donor cells (Fig. 4C). Monocytes accounted for 8.0% of recovered cells in C5-CD4 tumors, compared with 0.3% and 0.5% in CTRL-CD4 and sg*Il13*C5-CD4 tumors, respectively. Macrophages accounted for 4.1%, 1.2%, and 1.5% of recovered cells in the corresponding groups. Neutrophils and basophils each represented approximately 4 -5% of recovered cells in C5-CD4 tumors but 0.2% or less in the other groups (Fig. 5A).

**Figure 5.**
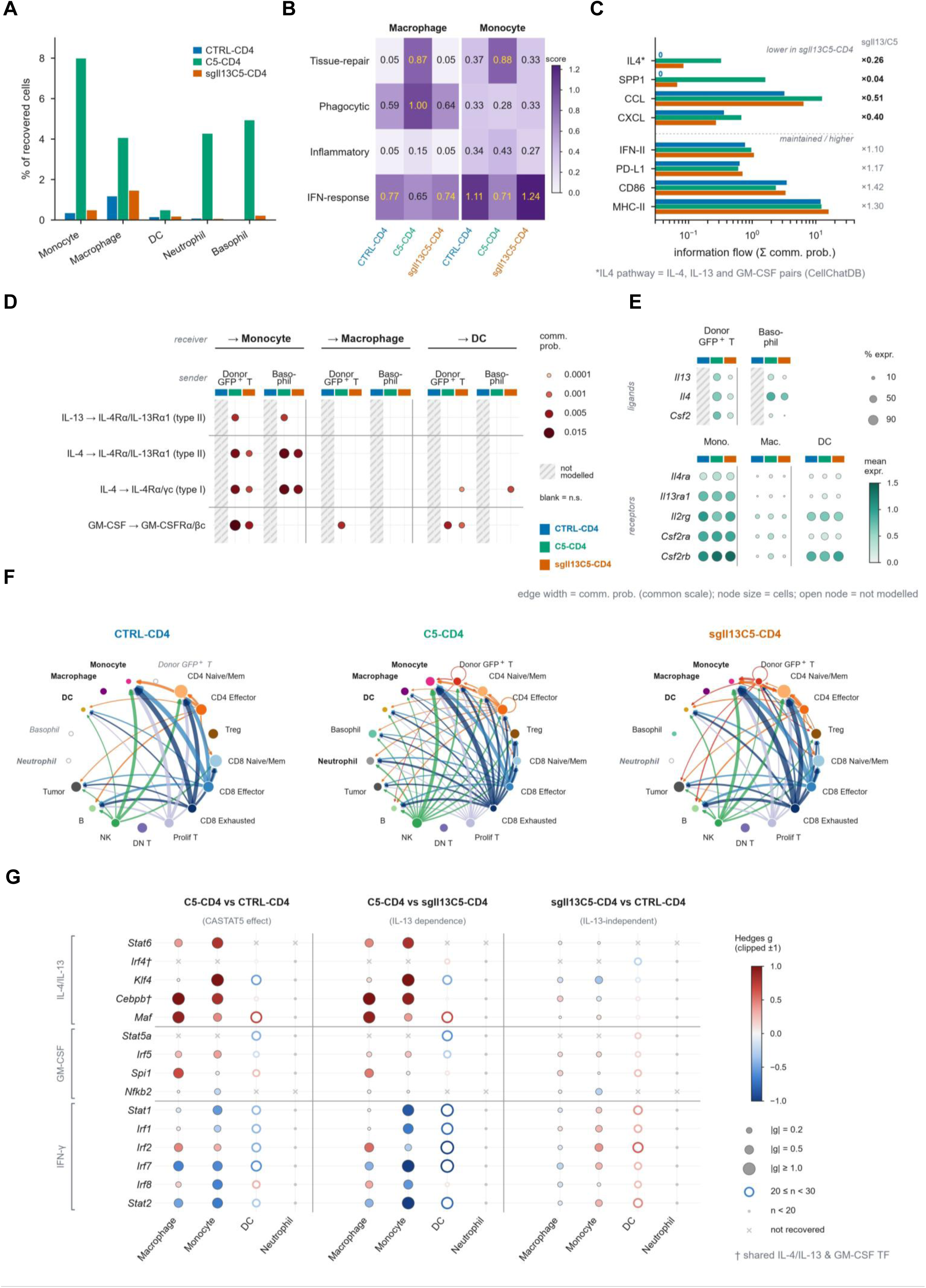
C5-CD4 T cell-derived IL13 regulates the abundance and phenotype of tumor-infiltrating myeloid cells. **A.** Relative abundance of major myeloid populations, including monocytes, macrophages, dendritic cells (DCs), neutrophils, and basophils, across treatment groups. **B.** Heatmap showing mean module scores for tissue-repair, phagocytic, inflammatory, and interferon (IFN) response programs in monocytes and macrophages. **C.** CellChat-inferred information flow across selected signaling pathways, calculated as the sum of communication probabilities for significant ligand–receptor interactions (permutation P < 0.05). Values are displayed on a logarithmic scale, with *sgIl13*C5-CD4/C5-CD4 ratios indicated on the right. The CellChatDB IL4 pathway includes IL-4, IL-13, and GM-CSF-associated signaling interactions. **D.** Predicted IL-13, IL-4, and GM-CSF ligand–receptor interactions from donor GFP⁺ T cells (C5-CD4 versus *sgIl3*C5-CD4) and basophils to monocytes, macrophages, and DCs. Dot size and color indicate communication probability; only significant interactions are shown. **E.** Dot plots showing expression of *Il13*, *Il4*, and *Csf2* in sender populations and their corresponding receptor genes (*Il4ra*, *Il13ra1*, *Il2rg*, *Csf2ra*, and *Csf2rb*) in myeloid populations. Dot size indicates the percentage of cells expressing each gene, and color represents mean log-normalized expression. Hatched columns in D and E indicate populations absent from the corresponding CellChat model. **F.** Circle plots showing CellChat-inferred IFN-γ–IFNGR1/IFNGR2 signaling networks across treatment groups. Edge width represents communication probability on a shared scale, and edge color identifies sender populations. Node size indicates cell abundance, with open nodes representing populations excluded from the model. Myeloid populations are highlighted in bold. **G.** Pairwise comparisons of pySCENIC regulon activity associated with IL-4/IL-13 (STAT6), GM-CSF (STAT5/NF-κB), and IFN-γ (STAT1/IRF) signaling in monocytes, macrophages, DCs, and neutrophils. AUCell scores were standardized within the corresponding regulon model, and differences were quantified using Hedges’ g. Dot size indicates effect magnitude, and color represents the direction of change, with positive values indicating higher activity in the first group. Open circles indicate limited cell representation (20-29 cells); gray dots indicate fewer than 20 cells; and crosses indicate regulons not recovered in the corresponding model. † indicates transcription factors shared by the IL-4/IL-13 and GM-CSF pathways. For CellChat analysis, populations with fewer than 20 cells were excluded from the corresponding group. CellChat P values were derived from interaction-level permutation tests.

In addition to differences in cellular composition, monocytes and macrophages from C5- CD4 tumors exhibited higher tissue-repair-associated program scores. Scores for macrophages and monocytes were 0.87 and 0.88, respectively, compared with 0.05 and 0.37 in CTRL-CD4 tumors and 0.05 and 0.33 in sg*Il13*C5-CD4 tumors. Conversely, the monocyte IFN-response program score was lower in C5-CD4 tumors than in CTRL-CD4 and sg*Il13*C5-CD4 tumors (0.71 versus 1.11 and 1.24, respectively; Fig. 5B). These findings indicate that the increased representation of myeloid cells in C5-CD4 tumors coincided with a shift toward tissue-repair-associated transcriptional programs and reduced IFN-response signatures.

To explore potential mechanisms underlying these changes, we performed CellChat analysis to infer intercellular communication. C5-CD4 tumors exhibited increased inferred information flow through the CellChatDB-defined IL4 pathway, which encompasses IL4-, IL13-, and GM-CSF-associated interactions in this analysis, as well as through the SPP1, CCL, and CXCL pathways. These signals were attenuated following donor *Il13* disruption, with sg*Il13*C5-CD4/C5-CD4 information-flow ratios of 0.26, 0.04, 0.51, and 0.40, respectively. In contrast, inferred IFN-II, PD-L1, MHC-II, and CD86 signaling was maintained or increased after *Il13* disruption, with corresponding ratios ranging from 1.10 to1.42 (Fig. 5C).

At the ligand–receptor level, CellChat predicted IL13 signaling through the type II IL4 receptor complex (IL4Rα/IL13Rα1) predominantly targeting monocytes in C5-CD4 tumors, with donor GFP+ T cells and basophils identified as potential ligand sources. Donor-derived IL4 and GM-CSF signaling to monocytes was also reduced approximately 4- to 6-fold following *Il13* disruption, whereas basophil-derived IL4 signaling was partially preserved (Fig. 5D). These inferred communication patterns coincided with lower expression of *Il13, Il4,* and *Csf2* in sg*Il13*C5-CD4 donor T cells. Mean log-normalized expression values were 0.10, 0.24, and 0.28, respectively, compared with 0.54, 0.63, and 0.71 in C5-CD4 donor cells. Basophil *Il13* expression was also lower in sg*Il13*C5-CD4 tumors (0.07 versus 0.30). Notably, monocyte expression of *Il4ra* and *Il13ra1* was comparable across groups, whereas both receptor-subunit transcripts were expressed at low levels in macrophages (Fig. 5E).

CellChat analysis further predicted IFN-γ signaling, primarily from CD8^+^ T cells and NK cells, toward monocytes and dendritic cells across all groups, whereas predicted signaling to macrophages was detected only in C5-CD4 tumors (Fig. 5F). Consistent with the inferred signaling patterns, regulon analysis revealed differences in transcriptional programs associated with IL4/IL13–STAT6 and IFN-γ–STAT1 signaling in monocytes. Compared with sg*Il13*C5-CD4 monocytes, C5-CD4 monocytes exhibited higher activity of the Stat6, Klf4, and Cebpb regulons (Hedges’ *g* = 0.81, 1.45, and 0.92, respectively) and lower activity of the Stat1, Stat2, Irf7, and Irf1 regulons (*g* = −0.86, −1.17, −1.08, and −0.69, respectively). By contrast, regulon activities differed only modestly between sg*Il13*C5-CD4 and CTRL-CD4 monocytes (|*g*| ≤ 0.41). Macrophages exhibited a similar, although less pronounced pattern, with higher Cebpb, Maf, and Spi1 regulon activity and lower Irf7 and Stat2 activity in C5-CD4 tumors (Fig. 5G).

Collectively, these analyses associate donor-derived IL13 with increased representation of monocytes, macrophages, neutrophils, and basophils, together with tissue-repair-associated myeloid transcriptional programs and inferred intercellular communication patterns consistent with enhanced IL4/IL13-STAT6-associated activity relative to IFN-response programs. The attenuation of these changes following *Il13* disruption in donor C5-CD4 T cells supports a role for IL13 in shaping the composition and transcriptional states of the tumor-infiltrating myeloid compartment.

### IL13 contributes to the efficacy of CASTAT5-transduced CD19CAR-T cells

Recent studies have reported that CD19-targeted CAR-T infusion products exhibiting a Th2-associated transcriptional program are linked to more favorable clinical outcomes, and that higher levels of type 2 cytokines, including IL4, IL5 and IL13, are detected in the sera of long-term responders(*35*). These observations raised the possibility that Th2-associated cytokines may contribute to the sustained therapeutic activity of CD19CAR-T cells. We previously demonstrated that expression of CASTAT5 enhances the antitumor activity of CD19CAR-T cells in the murine A20 B-cell lymphoma model(*22*). Given the prominent IL13 production by CASTAT5-conditioned CD4^+^ T cells and the association between Th2 features and favorable CAR-T outcomes, we therefore asked whether IL13 contributes to the enhanced therapeutic activity of CASTAT5-expressing CD19CAR-T cells. To address this question, mice bearing established A20 tumors were treated with regular CD19CAR-T cells, CASTAT5-CD19CAR-T (C5-CD19CAR) cells, or *Il13*-edited CASTAT5-CD19CAR-T (sg*Il13*C5-CD19CAR) cells (Fig. 6A). The three CAR-T products exhibited comparable CAR expression, as determined by Fab staining, and comparable CASTAT5 expression, as indicated by GFP expression (Supplemental Fig. 10A). Targeted disruption of the *IL13* locus in sg*Il13*C5-CD19CAR-T cells was confirmed by T7E1 restriction enzyme digestion (Supplemental Fig. 10B). At the dose used in this study, the regular CD19CAR- T cells produced only modest tumor growth delay, and all mice (8 of 8) succumbed to tumor progression by day 40. In contrast, C5-CD19CAR-T cells markedly controlled tumor growth and induced durable tumor regression, with 8 of 9 mice surviving long term through day 50 (Fig. 6B- C). Notably, disruption of IL13 substantially impaired the therapeutic potency of C5-CD19CAR-T cells. Mice treated with sg*Il13*C5-CD19 CAR-T cells underwent earlier tumor relapse and had a marked reduction in long-term survival, with only 1 of 7 mice surviving through day 50. These results implicate IL13 as an important functional component of the enhanced antitumor activity conferred by CASTAT5 in CD19CAR-T cells, extending the requirement for IL13 to genetically engineered CAR-T cells.

**Figure 6.**
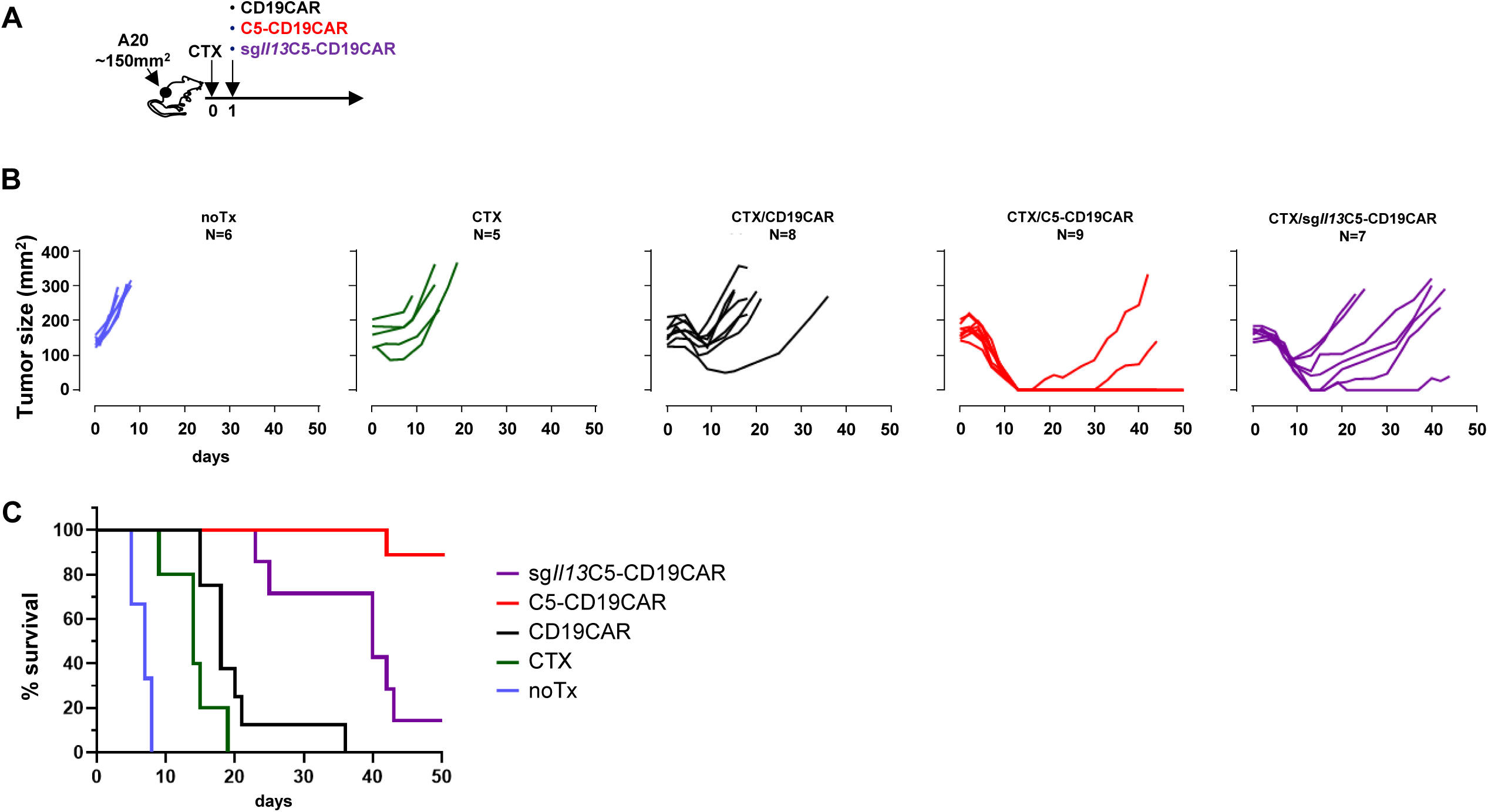
IL13 deficiency diminishes the efficacy of CASTAT5-modified CD19CAR-T cells. **A.** Schematic of the treatment groups and experimental design. **B**. Tumor growth curves for individual mice in each treatment group. **C**. Kaplan–Meier survival analysis of mice shown in B.

### IL13 expression in CD19CAR-T infusion products is associated with CAR-T persistence and patient outcomes

Bai et al. demonstrated that the Th2 gene signature (*IL4*, *IL5*, *IL13*, and *GATA3*) in CAR T infusion products was associated with ultra-long-term B cell aplasia and durable remission in pediatric B-ALL(*35*). To assess the association between type-2 functionality and *STAT5A* expression, we reanalyzed the single-cell infusion-product dataset reported by Bai et al. (GSE262072). In the discovery cohort (n = 42), products from patients with ultra-long-term B-cell aplasia (BCA-L, n = 5; 82–106 months) showed significantly higher Th2 signatures in both CD4⁺ and CD8⁺ CAR T cells than products from the remaining patients (2.3-fold, *P* < 0.001), with the difference driven primarily by *IL4* and *IL13* (Fig. 7A). Healthy-donor products showed type 2 cytokine expression levels comparable to those of BCA-L products. Interestingly, IL13 expression was similar between CD4⁺ and CD8^+^ T cells, whereas IL4 expression was lower in CD8^+^ T cells. Across the 42 discovery-cohort patients, the Th2 score and IL13 are nearly collinear (Pearson r = 0.962, P = 3×10⁻²⁴), indicating that *IL13* contributed substantially to the overall signature. *STAT5A* expression was also positively correlated with the average of Th2 signature genes (r = 0.44, *P* = 0.004), *IL4* (*r* = 0.39, *P* = 0.010), and *IL13* (*r* = 0.55, *P* = 0.0002), but not *with GATA3* (Fig. 7B).

**Figure 7.**
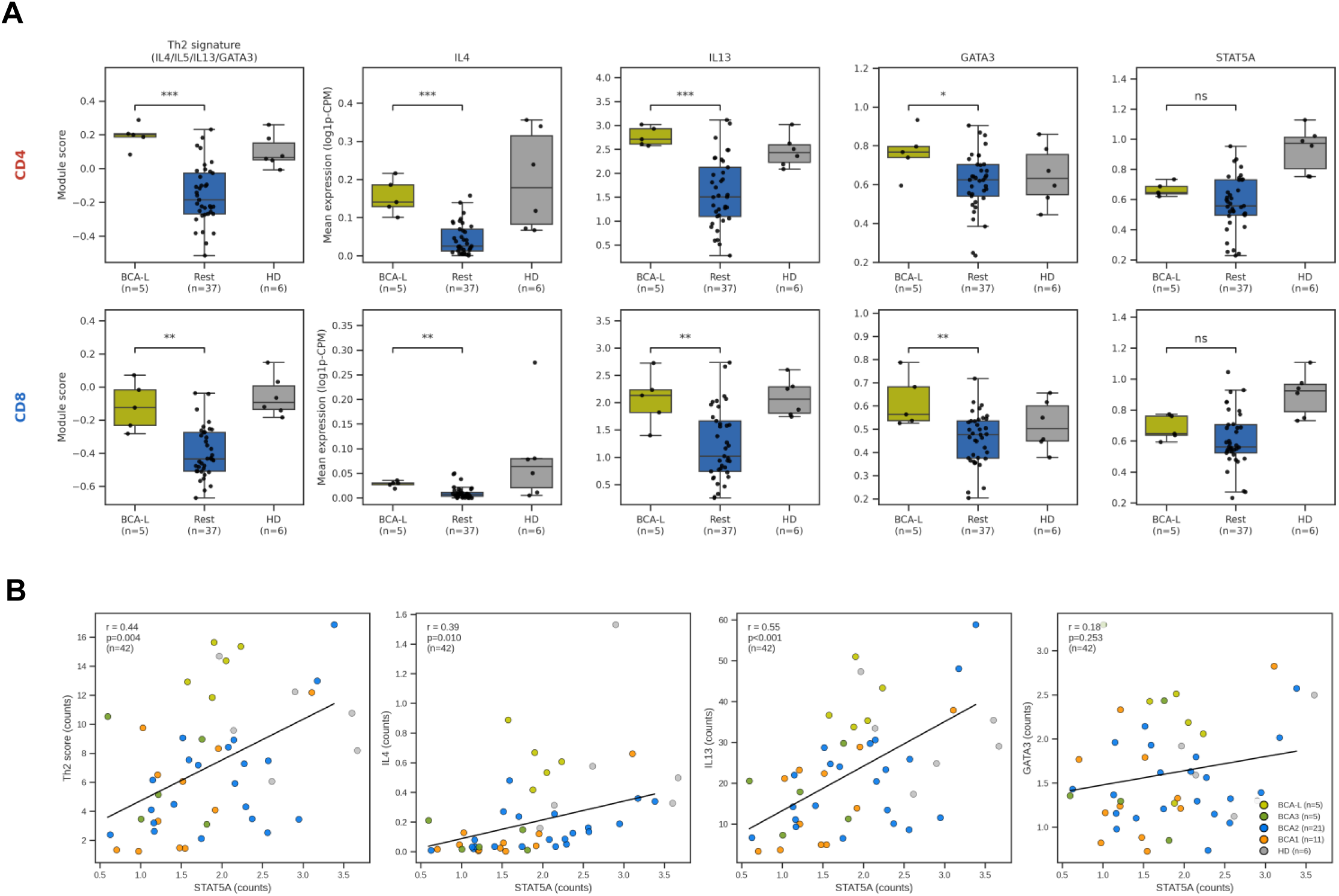
IL13 expression in CD19 CAR-T infusion products is associated with CAR-T persistence. **A.** Type-2 signature and gene expression in CD19-targeted CAR-T infusion products following in vitro antigenic stimulation. Data are from the discovery cohort (n = 42) in GSE262072, stratified by clinical persistence. CD4⁺ (top) and CD8⁺ (bottom) CAR-T cells were identified using CITE-seq antibody-derived tags. The Th2 signature (*IL4*, *IL5*, *IL13*, and *GATA3*) was calculated using Seurat AddModuleScore, and individual gene expression is shown as per-patient mean log1p-CPM values. Patients were classified as having ultra-long-term B-cell aplasia (BCA-L, n = 5) or as belonging to the remaining cohort (Rest, n = 37). Healthy donors (HD, n = 6) were included as references but excluded from statistical comparisons. Boxes indicate medians and interquartile ranges (IQRs); whiskers extend to 1.5 × IQR; dots represent individual patients. Two-sided Mann–Whitney U test; *, P < 0.05, **, P < 0.01, ***, P < 0.001. **B.** Correlation between *STAT5A* expression and the type-2 transcriptional signature across CD19CAR-T infusion products from the discovery cohort (n = 42). Per-patient mean *STAT5A* expression (x-axis) was plotted against the type 2 signature score and the expression of individual genes (*IL4, IL13,* and *GATA3*; y-axis). Individual gene expression values were calculated by pseudobulk analysis using raw counts per cell. Each dot represents an individual patient, colored by clinical BCA persistence group (BCA-L, BCA3, BCA2, and BCA1). Healthy donors are shown in gray for reference and are excluded from correlation analyses. Lines indicate least-squares linear regression, with Pearson correlation coefficients (r) and corresponding P values.

However, *STAT5A* expression was not associated with BCA-L status or BCA duration. These findings link *STAT5A* expression to type-2 cytokine programs in CD19CAR-T infusion products but do not establish a direct association between STAT5A expression and with clinical CAR-T persistence.

## Discussion

Although IL13 has traditionally been associated with tumor-promoting inflammation and evasion of immunosurveillance(*42*), our findings reveal an unexpected antitumor role for IL13 in the context of ACT using polyfunctional CD4^+^ T cells. Here we identify IL13 as a critical effector cytokine contributing to the durable antitumor activity of CASTAT5-conditioned polyfunctional CD4⁺ T cells (C5-CD4). Genetic disruption and antibody-mediated neutralization of IL13 impaired C5-CD4-mediated tumor control, while IL13 overexpression alone was insufficient to recapitulate the full therapeutic activity conferred by CASTAT5. These complementary findings identify IL13 as a functional component of the CASTAT5-induced effector program rather than simply a marker of Th2 differentiation, and provide genetic evidence that T cell-derived IL13 contributes to the efficacy of adoptively transferred polyfunctional CD4⁺ T cells. Our results further associate IL13 with donor T-cell persistence, remodeling of the tumor-associated myeloid compartment, and activation of endogenous tumor-reactive CD8⁺ T cells, suggesting that its therapeutic contribution extends beyond the transferred cells themselves.

In our model, IL13 is produced as part of a broader CASTAT5-driven polyfunctional program that includes IL4, GM-CSF, IFNγ, TNFα, and cytolytic activity. Epigenomic analyses revealed extensive chromatin remodeling by CASTAT5 at Th2 cytokine loci (Fig. 2), providing a potential regulatory basis for the prominent type 2 cytokine profile of C5-CD4 cells. Importantly, IL13 overexpression alone did not reproduce their full therapeutic activity (Fig. 3H), supporting a model in which IL13 operates within a coordinated effector network rather than acting as an autonomous antitumor factor. These findings underscore the importance of evaluating cytokine function in the context of the broader cellular program in which it is expressed.

An important observation is that IL13 disruption affected the abundance and persistence of transferred C5-CD4 cells. Despite retaining expression of several activation-associated genes and effector cytokines, IL13-deficient C5-CD4 T cells declined progressively after transfer and showed increased expression of inhibitory-receptor-associated genes, including *Tox*, *Ctla4, and Lag3*. These findings suggest that IL13 may help sustain donor cell fitness or persistence in the TME, although the underlying mechanism remains unresolved. Because fewer donor cells could by themselves lower systemic cytokine levels, myeloid recruitment and endogenous CD8⁺ T-cell priming, some of the downstream changes associated with IL13 loss may be secondary to impaired donor persistence. Consistent with this possibility, serum IFNγ, TNFα and GM-CSF were reduced even though sg*Il13*C5-CD4 T cells retained, or increased, their capacity to produce these cytokines on restimulation (Fig. 3C-D). Consistent with previous studies(*26, 42*), we found that both donor and endogenous T cells expressed little or no *Il13ra1*, suggesting that direct autocrine or paracrine effects of IL13 on T cells are less likely. Instead, IL13 may indirectly support T-cell persistence or function by modifying the surrounding immune microenvironment, potentially through IL13-responsive myeloid cells. This possibility contrasts with the reported direct effects of IL4 on T cells in ACT, where IL4 signaling can enhance donor T cell metabolic fitness and resistance to exhaustion(*43*). Interestingly, however, IL4 neutralization did not compromise the therapeutic efficacy of C5-CD4 cells in our model. One possible explanation is that the beneficial T cell-intrinsic effects of IL4 signaling, including enhanced metabolic fitness and resistance to exhaustion, are compensated by alternative pathways activated by CASTAT5, thereby diminishing the requirement for IL4. Thus, although IL13 and IL4 share aspects of type 2 immune signaling, they may contribute to T cell-mediated antitumor immunity through distinct cellular targets and mechanisms.

Our single-cell analyses revealed increased representation of tumor-infiltrating monocytes and macrophages expressing tissue-repair-associated transcriptional programs in C5-CD4-treated tumors (Fig. 5A). Notably, this myeloid cell accumulation was observed 7 days after T cell infusion, preceding the complete tumor regression typically observed 10–15 days after transfer in survival studies. Macrophages in C5-CD4-treated tumors also exhibited higher tissue-repair and phagocytic program scores than those in CTRL-CD4-treated tumors, while still retaining inflammatory and IFN-responsive programs (Fig. 5B). Monocytes from C5-CD4-treated tumors likewise showed higher tissue-repair and lower IFN-response program scores. We speculate that the timely accumulation of reparative myeloid cells may contribute to clearance of dying tumor cells and restoration of tissue homeostasis, potentially limiting prolonged inflammation that could otherwise undermine durable tumor control. However, the functional contributions of these cells to tumor regression remain to be established. Our findings associate donor-derived IL13 with accumulation of monocytes/macrophages exhibiting tissue-repair-associated transcriptional programs. Although these macrophage states share phenotypic features with alternatively activated macrophages and M2-like tumor-associated macrophages (TAMs), which are often associated with tumor immune evasion, their functional properties cannot be inferred from transcriptional markers alone. In C5-CD4-treated mice, the concurrent presence of abundant proinflammatory cytokines, including IFNγ, TNFα, and GM-CSF (Fig. 3D), may prevent tissue- repairing macrophages from adopting an immunosuppressive phenotype. The functional characterization of these tumor-infiltrating myeloid populations and the contributions of IL13 and other cytokines to their differentiation and activity warrant further investigation.

Residual A20 tumor cells in C5-CD4-treated mice showed a glycolytic and hypoxia- associated stress signature and reduced expression of mature B-cell identity genes, and bothfeatures were largely reversed after donor *Il13* disruption (Supplemental Fig. 9). Because A20 cells expressed little *Il13ra1* or *Il13ra2*, these changes are unlikely to reflect direct IL13 signaling in tumor cells. Type 2 immunity has been reported to restrain tumor growth through vascular remodeling that leads to tumor hypoxia and cancer cell death(*9*), and CD4⁺ T cells can damage tumor vasculature through IL-3 and TNFα(*12*). An indirect route of this kind, acting through IL13- responsive myeloid cells or the tumor vasculature, is therefore plausible and warrants further investigation.

The clinical relevance of our polyfunctional CD4⁺ T cell ACT model is supported by emerging evidence from CD19-targeted CAR-T therapies. Polyfunctional CD4⁺ CD19CAR-T cells in preinfusion products correlate with favorable clinical outcomes(*20*), and CD4⁺ CD19CAR-T cells constituted the predominant donor-cell population in two patients with remissions lasting more than 10 years(*44*). Bai et al. associated type 2 cytokine signatures in preinfusion products with prolonged B-cell aplasia and patient survival, and detected higher serum IL13 in long-term responders (*35*). Our reanalysis of this dataset confirms that the difference in type 2 signature is driven primarily by *IL4* and *IL13* (Fig. 7), although whether IL13 contributes directly to clinical efficacy remains unknown. Activated human CD19CAR-T cells already co-express type 1 and type 2 cytokines(*45*). Our finding that IL13 contributes to the efficacy of CASTAT5-expressing CD19CAR-T cells in mice raises the possibility that a similar mechanism operates in human CAR-T therapy.

Our study has several limitations. Many types of cancer express receptors for IL4 and IL13 and may therefore respond to these cytokines with enhanced growth or survival(*29, 42*). The therapeutic effects of type 2 cytokines produced by donor T cells may be contingent on the accompanying cytokine milieu. Further investigation is needed to define the conditions under which T-cell infusion products containing type 2 cytokine programs may be therapeutically beneficial or potentially detrimental, taking into account factors such as donor T cell cytokine profiles and tumor types. Another important consideration is the potential for increased treatment-related toxicity associated with highly polyfunctional T cells. In clinical CD19 CAR-T studies, a higher polyfunctionality strength index (PSI) in preinfusion products has been associated with enhanced antitumor efficacy, but also with an increased risk of grade ≥3 cytokine release syndrome and neurotoxicity(*20*). Murine models have limited capacity to predict toxicities associated with human ACT. Whether the coordinated cytokine profile of CASTAT5-conditioned polyfunctional T cells provides a more favorable balance between antitumor efficacy and treatment-related toxicity remains an important question for future investigation.

In summary, our findings reveal a context-dependent antitumor role for IL13 produced by CASTAT5-conditioned polyfunctional CD4^+^ T cells. IL13 contributes to optimal therapeutic activity, is associated with remodeling of the tumor-associated myeloid compartment, and supports broader endogenous antitumor immune responses. The extension of this requirement to CASTAT5-engineered CD19CAR-T cells highlights the potential relevance of IL13-associated programs to engineered T-cell therapy. More broadly, this work illustrates that the functional effects of individual cytokines depend on the cellular context and the immune networks in which they operate.

### Material and Method Experimental mice

BALB/c mice (Thy1.2^+/+^) of 4 to 6 weeks of age were purchased from Charles River. 6.5 TCR-Tg mice on a BALB/c (Thy1.1^+/+^) background expressing an αβTCR specific for amino acids 110– 120 from influenza HA presented by MHC class II molecule IE^d^ were described previously(*46*). All mice were housed under specific pathogen-free (SPF) conditions by Laboratory Animal Services of Augusta University. All animal experiments were approved by the Institutional Animal Care and Use Committee (IACUC) of Augusta University.

### Antibodies and reagents

The following anti-mouse antibodies, including CD4 (GK1.5), CD3 (17A2), CD90.1 (OX-7), PD-1 (RMP1-14), CD44 (IM7), CD11C (N418), CD45.1 (A20), anti-IL-2 (JES6-5H4), anti-TNF-α (MP6-XT22), anti-IFN-γ (XMG1.2), anti-IL4 (11B11), anti-IL-9 (RM9A4), anti-GM-CSF (MP1-22E9), anti-Granzyme B (GB11), anti-GATA3 (16E10A23) and anti-Stat5 Phospho (Tyr694) (A17016B), and anti-human antibodies, including CD4 (OKT4), CD8a (HIT8a) and Bcl2 (100), were purchased from Biolegend. Anti-mouse CD25 (PC61), EZH2 (11/EZH2) and Ki-67 antibodies were purchased from BD Biosciences. Anti-mouse antibodies IL13 (eBio13A) and Bcl-2 (Bcl2/100) were purchased from eBioscience. Alexa Fluor 488-conjugated goat anti-rabbit IgG antibody was purchased from Thermo Fisher Scientific. FITC-conjugated polyclonal mouse anti-rat F(ab)2 antibodies and FITC-conjugated polyclonal mouse IgG antibodies were purchased from Jackson ImmunoResearch. IL13 neutralizing antibody anti-mIL13-mIgG1 (clone 8H8) was purchased from Invivogen. IL4 neutralizing antibody (clone 11B11) was purchased from BioXCell. Intracellular fixation and permeabilization solution kit was purchased from BD. Dynabeads mouse T-activator, Dynabeads human T-activator, and transcription factor staining buffer set were purchased from Thermo Fisher Scientific. Cyclophosphamide monohydrate (CTX) was purchased from Tokyo Chemical Industry. Retronectin was purchased from Takara. Lipofectamine 2000 was purchased from Invitrogen.

### Tumor cell lines

The murine B-cell lymphoma cell line A20 was obtained from American Type Culture Collection (ATCC). The hemagglutinin-expressing A20 tumor cell line (A20HA) was generated as described previously(*19*). Tumor cells were cultured in RPMI 1640 (HyClone Laboratories) supplemented with 10% fetal bovine serum albumin (FBS), 1% penicillin/streptomycin (HyClone Laboratories), 1% non-essential amino acids, and 1% glutamine (Corning) at 37 °C in a 5% CO2 incubator.

### Animal tumor models and treatments

Tumor cells were subcutaneously injected into the right flank of mice (2.5 x 10^6^ tumor cells/mouse in 50 ul PBS). Tumor sizes were measured by a caliper every other day and expressed as the product of two perpendicular diameters in square millimeters. When tumor sizes reached the desired sizes, mice were randomly assigned into groups to receive the specified treatments. CTX was dissolved in PBS and i.p. injected into mice at a dose of 150 mg/kg one day before T cell transfer. For adoptive T-cell transfer, in vitro cultured T cells were harvested, washed twice with PBS, and injected into each recipient via the tail vein. For *in vivo* neutralization of IL4 or IL13, mice received i.p. injections of neutralizing anti-IL4 (200 µg per dose) or anti-IL13 (100 µg per dose) antibodies twice weekly for a total of six doses, starting immediately after T-cell infusion.

### Guide RNA design, RNP complex formation, and nucleofection of activated T cells

Targeted gene editing was performed following the procedures reported by Seki et al(*41*), with modifications using the Alt-R CRISPR-Cas9 system from Integrated DNA Technologies (IDT). To ensure targeted deletion of the *Il13* gene, predesigned guide RNAs targeting exons 1 and 3 of *Il13* on mouse chromosome 11 were selected based on their predicted high on-target activity and low off-target activity. Predesigned CRISPR RNAs (crRNAs), trans-activating CRISPR RNAs (tracrRNAs), and Cas9 protein were purchased from IDT. Scramble crRNAs #1 and #2 were included as controls. All crRNA sequences are provided in Supplemental Table I. For RNP complex formation, crRNAs were reconstituted to 100 μM in nuclease-free duplex buffer and mixed with tracrRNA at equal concentrations. The RNA components were annealed by heating at 95°C for 5 min, followed by gradual cooling to room temperature over 30–60 min. The resulting crRNA–tracrRNA duplexes were mixed with Cas9 protein and incubated at room temperature for at least 10 min to form RNP complexes. T cells (1-3 × 10⁶) were pelleted in 1.5-mL microcentrifuge tubes and resuspended in 20 μL of P3 Primary Cell Nucleofector Solution (Lonza). The cell suspension was mixed with the preassembled RNP complexes and transferred to nucleofection cuvettes using the Amaxa P3 Primary Cell 4D-Nucleofector Kit S (Lonza). Nucleofection was performed using the 4D-Nucleofector system with program CM137 for activated murine T cells. Following nucleofection, cells were transferred to 24-well recovery plates containing 80 μL of prewarmed T-cell medium and cultured with recombinant IL-2 and IL-7 for 4 h before subsequent manipulation.

### Gene-editing validation by T7 endonuclease I (T7E1) digestion and Nanopore sequencing

Genomic DNA was isolated from control and gene-edited CD4⁺ T cells using the DNeasy Blood & Tissue Kit (Qiagen). For Nanopore sequencing, the target genomic region was amplified by PCR using Q5 Hot Start High-Fidelity 2X Master Mix (New England Biolabs), per the manufacturer’s instructions. For T7E1 digestion, the target region was amplified using DreamTaq PCR Master Mix (2X) (Thermo Scientific), following the manufacturer’s protocol. PCR products were then subjected to heteroduplex formation and digestion with T7 Endonuclease I (New England Biolabs), per the manufacturer’s instructions. The sequences of the forward and reverse primers are provided in Supplemental Table I.

### Mouse T cell retroviral transduction and adoptive transfer

The murine stem cell virus (MSCV) retroviral vectors encoding CASTAT5 and GFP or murine IL13 and mCherry, each separated by a P2A sequence, were purchased from VectorBuilder Inc. The MSGV-1D3-28Z.1-3 retroviral vector encoding an anti-CD19 chimeric antigen receptor (CAR), in which the first and third immunoreceptor tyrosine-based activation motifs (ITAMs) of the CD3ζ chain were inactivated to reduce apoptosis, was provided by Dr. James N. Kochenderfer(*47*). For CASTAT5 transduction, Phoenix-Eco cells (ATCC) were transfected with the CASTAT5-GFP retroviral vector and the pcL-Eco packaging plasmid using Lipofectamine 2000 (Invitrogen), according to the manufacturer’s instructions. Viral supernatants were collected 48, 72, and 96 h after transfection. Mouse CD4^+^ T-cell transduction was performed as previously described(*22*). Briefly, HA-specific CD4^+^ T cells were activated overnight using CD3/CD28 T-cell activation Dynabeads (Invitrogen) and then transduced twice with CASTAT5 retroviral supernatant on consecutive days. Cells were maintained in medium supplemented with hIL-2 (30 U/mL) throughout the transduction procedure. Twenty-four hours after the second transduction, cells were harvested, and the Dynabeads were removed by magnetic separation. Cells were then sorted using a Bigfoot Cell Sorter into GFP^+^ (CASTAT5-transduced) and GFP^−^ (untransduced control) populations before adoptive transfer into recipient mice. To prepare IL13-deficient CASTAT5-CD4 T cells (sg*Il13*C5-CD4), the nucleofection procedures described above were performed on day 2 before the first viral transduction.

### Sample preparation and flow cytometry analysis

Spleen and tumor samples were processed into single-cell suspensions. Peripheral blood was collected via tail vein bleeding. Red blood cells in samples were lysed with ACK lysing buffer. For surface molecule detection, cells were stained with fluorochrome-conjugated antibodies for 15 minutes at room temperature in the dark. To detect intranuclear molecules, the transcription factor staining buffer set (eBioscience) was used following the manufacturer’s instructions. For intracellular cytokine staining, samples were stimulated with Leukocyte Activation Cocktail containing PMA/ionomycin in the presence of GolgiStop (BD Biosciences) for 4 hours at 37 °C. Cells were harvested and surface stained, followed by cytokine staining with intracellular fixation and permeabilization kit (BD Biosciences). To detect phosphorylated Stat5 (pStat5) in T cells, cells were fixed in 2% formaldehyde followed by permeabilization with ice-cold methanol. After washing with PBS, cells were stained with pStat5 antibody at room temperature for 20 minutes before washing and flow cytometry analysis. All FACS data were acquired on an Attune NxT (Thermo Fisher) or NovoCyte Quanteon (Agilent) Flow Cytometer and analyzed using FlowJo software (Tree Star).

### Luminex quantitation of cytokines and chemokines in mouse serum

Detection of cytokines in serum was performed using Mouse Procarta 26-Plex Cytokine & Chemokine Panel 1 (Thermo, Cat# EPX260-26088-901) according to manufacturer’s instructions. In brief, beads were added to a 96-well plate and washed in Bio-Plex Pro Wash Station (Bio-Rad Cat# 30034376). Samples were diluted 2-fold with Universal Assay Buffer, added to the plate and incubated with magnetic bead-conjugated capture antibodies overnight at 4°C on a plate shaker (350 rpm). After washing the plate, detection antibodies were added to each well and the plate was incubated on a shaker (600 rpm) for 30min at room temperature. Finally, after washing, SA- PE was added and incubated for 30 min. Samples were acquired on a Luminex FLEXMAP 3D instrument (Thermo, Cat# APX1342). Standard curves and protein concentration were calculated using Quantist Luminex Data Analysis Software (Bio-Techne).

### Nanopore long-read sequencing analysis of the edited genomic locus

To detect CRISPR/Cas9-induced gene-editing events, PCR primers were designed to flank the sgRNA target site, enabling amplification of the edited genomic regions. Purified PCR products were prepared for sequencing with the Oxford Nanopore Ligation Sequencing Kit V14 (SQK-LSK114) following the manufacturer’s protocol. The resulting libraries were sequenced on an Oxford Nanopore MinION device (Product code: PRO-SEQ 002). Basecalling was performed with the Dorado v8 basecaller, generating high-quality sequence reads for each amplicon. The long-read sequencing data were then visualized and analyzed in the Integrative Genomics Viewer (IGV) to inspect editing outcomes, including indel patterns and structural alterations at the target locus.

### CUT&Tag library preparation and analysis

CUT&Tag libraries were prepared with the CUT&Tag-IT Assay Kit (Active Motif, 53160), following the manufacturer’s protocol with minor modifications. Donor C5-CD4 T cells and endogenous CD4⁺ T cells were isolated from mouse spleens via FACS sorting based on the Thy1.1 congenic marker at the indicated time points. 200,000 -350,000 cells were used per reaction. Samples were incubated overnight with antibodies against H3K4me3 (39060), H3K27ac (39034), or H3K27me3 (39055), all from Active Motif. Libraries were amplified for 10–22 cycles. Libraries were sequenced by MedGenome or the Georgia Cancer Center Genomics Core, with a reported target of 5-10 million paired-end reads per sample. CUT&Tag reads were processed on Galaxy. Adapter and quality trimming used Trim Galore! v0.6.10, followed by alignment to mm10 with Bowtie2 v2.5.321. PCR duplicates were removed with MarkDuplicates v3.1.1.0. Mitochondrial alignments and reads overlapping ENCODE blacklist v2 regions were excluded. Coverage tracks were generated with deepTools. Peak calling used MACS2 callpeak with IgG normalization. Three biological replicates per mark and condition were used. Differential CUT&Tag peaks were identified with DiffBind v3.12.0 using its DESeq2 engine(*48*). Quantification used 400-bp intervals centered on peak summits. An adjusted p-value below 0.05 defined significant changes. Peaks were annotated with ChIPseeker(*49*), and genomic overlaps were identified with GenomicRanges findOverlaps. For display, replicate bigWig tracks were averaged per mark and condition, and each mark was drawn on a common y-axis for both conditions, capped at the 99.5th percentile of the pooled signal in the displayed window.

### Hi-C library preparation

Hi-C libraries were prepared using the Arima Genome Wide HiC+ Kit (A510008) and the library preparation kit (A303011). Sorted inputs comprised 1.6 million endogenous CD4^+^ T cells and 3.4 million C5-CD4 T cells, isolated from mouse spleens by FACS using the Thy1.1 congenic marker at the specified time points. Reads were processed with Juicer v1.6(*50*) against mm10 to generate .hic contact maps. Chromatin loops were called with HiCCUPS at 10-kb resolution, and the enriched-pixel calls were classified by anchor overlap: a call in one condition was scored as shared if the other condition had a call within ±1 bin at both anchors, and otherwise as condition-specific (C5-CD4-gained, 9,982 calls; shared, 17,019; Endo. CD4-only, 2,279; shared in Endo. CD4, 12,677). At 17,019 shared CASTAT5-reference coordinates, log₂ contact-count ratios were corrected for total valid pairs: 425,041,512 in CASTAT5 and 311,060,166 in controls, a ratio of 1.366. Contact matrices were converted to cooler format and analyzed with cooler and cooltools. For display, SCALE-normalized 10-kb contact matrices were exported to cooler format and rendered with cooler/matplotlib as rotated triangle maps for each condition, on a common color scale capped at the 99th percentile of off-diagonal values within 50 bins (500 kb) of the diagonal, together with a log2 contact-ratio map, log2[(C5-CD4 + 1)/(Endo. CD4 + 1)], computed bin by bin.

### Analysis of the external STAT5 ChIP-seq reference

Public STAT5 ChIP-seq data from GSE207265, which profiled IL2-stimulated mouse CD4⁺ effector T cells in a separate study(*40*), provide reference occupancy annotations. They do not measure differential STAT5A binding in the CASTAT5 and control samples analyzed here. Reference bigWig tracks were converted from mm9 to mm10 using CrossMap Wig (Galaxy v0.7.0) and the UCSC mm9ToMm10 chain. STAT5 peaks were annotated with ChIPseeker(*49*).

### Integrated analysis of Hi-C, CUT&Tag of the Th2 and Gata3 regulatory regions

For the Th2 cytokine locus (chr11:53.4–54.4 Mb; contact maps 53.45–54.32 Mb) and the Gata3 locus (chr2:9.2–10.4 Mb), loops were counted only if both anchors were within the displayed window. C5-CD4-gained and Endo. CD4-only loops are individual 10kb HiCCUPS calls; shared loops were collapsed into a single contact by merging shared calls from both conditions (C5-CD4 and Endo. CD4) that were within ±1 bin of each other at both anchors, resulting in one shared contact per locus (Th2: 15 gained, 3 Endo. CD4-only, 5 shared; Gata3: 13, 2, 4). In the browser view, loop arcs, the STAT5 reference track, and the replicate-averaged H3K4me3, H3K27ac, and H3K27me3 signals are shown alongside UCSC refGene (mm10) gene models. The STAT5 reference track is the mean of two ChIP-seq replicate bigWigs, and the STAT5 peaks are the union of the two replicate peak sets, merged with GenomicRanges::reduce (34,289 peaks). Loop anchors were annotated with STAT5 peaks and with differential CUT&Tag peaks (DESeq2 tables per mark) overlapping the 10kb anchor ±5kb; a peak was scored as gained or lost at p < 0.05 and |log2 fold change| ≥ 0.5. As a control for the higher sequencing depth of the C5-CD4 library, the raw 10kb cis contacts were extracted from both .hic files; the C5-CD4 contacts were binomially thinned to the Endo. CD4 total (311.05 million); both maps were then rebuilt (juicer_tools 2.20.00, SCALE normalization), and HiCCUPS was re-run at 10kb with default parameters; a loop was considered to pass if it was still called in the thinned C5-CD4 map and still absent from the rebuilt Endo. CD4 map under the same ±1 bin rule. At matched depth, 13 of the 15 C5-CD4-gained loops at the Th2 locus and 6 of the 13 at the Gata3 locus were retained; all of the Endo. CD4-only loops and all of the shared contacts were retained.

### scRNA-seq analysis of tumor-infiltrating immune cells

Single-cell RNA-seq was performed on CD45.2^+^ cells, which were FACS-sorted from A20HA tumors resected from mice that received CTRL-CD4, C5-CD4, or sg*Il13*C5-CD4 T cells. For each treatment group, tumors were collected from three separate mice, and equal numbers of sorted cells from the three samples were pooled to generate a single composite specimen per group. Libraries were prepared using the 10x Genomics Chromium Single Cell 3’ v3.1 chemistry, sequenced on an Illumina NovaSeq 6000, and processed with Cell Ranger v8 against the mouse (mm10) reference, including the EGFP sequence. In Seurat (v5.4)(*51*), cells with ≤300 detected genes or ≥10% mitochondrial reads were removed, and doublets were excluded using DoubletFinder (v2.0.4), leaving 45,097 cells (CTRL-CD4, 18,281; C5-CD4, 10,841; sgIl13C5- CD4, 15,975). Data were log-normalized, integrated via canonical correlation analysis, reduced by PCA and UMAP, clustered by shared nearest neighbor modularity optimization, and annotated using cluster markers identified by FindMarkers. Donor cells were defined as those with ≥1 GFP transcript; host T cells were defined as GFP-negative T cells. All downstream analysis was performed in Python using scanpy, including gene signature scores (score_genes, equivalent to Seurat AddModuleScore) and re-clustering of the tumor cells. Tumor cells were identified using InferCNV(*52*). Cell-cell communication was inferred using CellChat(*53*) (v2.2.0.9001; CellChatDB.mouse; interactions retained at permutation p < 0.05), and transcription factor regulon activity was assessed using pySCENIC(*54*). Because each arm was a single pooled specimen, between-arm comparisons are purely descriptive. Expression changes are reported as log2 ratios of mean normalized expression; proportions are reported as percentages of cells (each with 95% intervals from cell resampling (bootstrap or Wilson score)); and regulon differences are reported as Hedges g, a measure of effect size. The reported intervals are from cell resampling only and do not represent between-animal variability.

### Previously generated datasets and the external Stat5 reference

ATAC-seq and bulk RNA-seq were obtained from GSE15224214. Public Stat5 ChIP-seq data were obtained from GSE207265, which profiled IL-2-stimulated mouse CD4 effector T cells in a separate study19. These data provide reference occupancy annotations; they do not measure differential Stat5A binding in the CASTAT5 and control samples analyzed here. Reference bigWig tracks were converted from mm9 to mm10 using CrossMap Wig (Galaxy v0.7.0) and the UCSC mm9ToMm10 chain. Stat5 peaks were annotated with ChIPseeker20.

### Reanalysis of human CD19CAR-T dataset

The processed single-cell Seurat object for the CTL019 infusion-product atlas (GSE262072), which contains 695,819 high-quality CAR T cells from 82 pediatric B-ALL patients and 6 healthy donors (basal and CD19-stimulated conditions), was downloaded from the Gene Expression Omnibus (GEO). Because the normalized data layer of the deposited object was identical to the raw counts, all analyses were performed on raw counts, with log1p-transformed counts-per- 10,000 (log1p-CPM) computed from counts as a normalization-robustness check. Per-patient clinical annotations (response, BCA group, BCA duration, relapse status) were obtained from the supplementary patient-demographics table of Bai et al. BCA groups were as defined in the original study by B cell aplasia duration: in the discovery cohort (trial NCT01626495, n = 42), BCA-L (n = 5; 82-106 months, ongoing), BCA3 (n = 5), BCA2 (n = 21) and BCA1 (n = 11; < 3 months); in the validation cohort (trial NCT02906371, n = 40), BCA-O (n = 11; 48-72 months, ongoing), BCA3 (n = 6), BCA2 (n = 17) and BCA1 (n = 6). Analyses used CD19-stimulated cells (12 h co-culture with CD19-expressing 3T3 feeder cells; CAR+ cells enriched with anti-FMC63-PE antibody and anti-PE magnetic beads). CD4+ and CD8+ cells were classified from CITE-seq antibody-derived tags (ADT): CD8+ if log1p(ADT-CD8) > 4.261 and log1p(ADT-CD4) -log1p(ADT-CD8) <= 0.5; CD4+ if log1p(ADT-CD4) - log1p(ADT-CD8) > 0.5 and log1p(ADT-CD8) <= 4.261; remaining cells were excluded as ambiguous. Gene expression was aggregated as per-patient mean raw counts within each subset (all stimulated cells for correlation analyses). The Th2 score was defined as the per- patient mean of IL4, IL5, IL13, and GATA3. Group comparisons used two-sided Mann-Whitney U tests (BCA-L, n = 5, vs the remaining discovery-cohort patients, n = 37; replicated as BCA-O, n = 11, vs the remaining validation-cohort patients, n = 27-28). Healthy donors (n = 6) were plotted as a reference group and were not included in statistical tests. Correlations across the 42 discovery-cohort patients used Pearson’s r; association with BCA duration used Spearman’s rho; differences between CD4+ and CD8+ compartment correlations were assessed with a patient-level bootstrap (10,000 resamples). Analyses were performed in Python (pandas, SciPy, seaborn).

### Statistical analysis

Statistical analyses were performed using GraphPad Prism 9.0 (GraphPad software under license by Augusta University). For comparison between two groups, the statistical significance of the results was determined using an unpaired two-tailed Student’s *t-*test. One-way ANOVA was used to determine statistical differences among three or more groups. For survival analysis, we employed the log-rank (Mantel-Cox) test.

## Acknowledgments

We thank the Flow and Mass Cytometry Core Facility at Georgia Cancer Center (RRID: SCR_025747) for maintaining and ensuring the smooth operation of the flow cytometers. We thank Rebekah Tritz and David Hansen for their assistance with cell sorting using the Bigfoot Spectral Cell Sorter. We acknowledge the support and contribution of the Integrated Genomics Core Shared Resources (RRID: SCR_026483) and the Immune Monitoring Core (RRID: SCR_026590) at the Georgia Cancer Center of Augusta University.

Author Contributions

ODO conducted experiments and analyzed data; MKI, XW, YY, AB, LP, CB, MYG, ZCD and RP assisted with experiments and data analysis; HS and GZ conceived and designed the study, conducted experiments, interpreted the results, and wrote the manuscript.

---

**Supplemental Figure 1.**
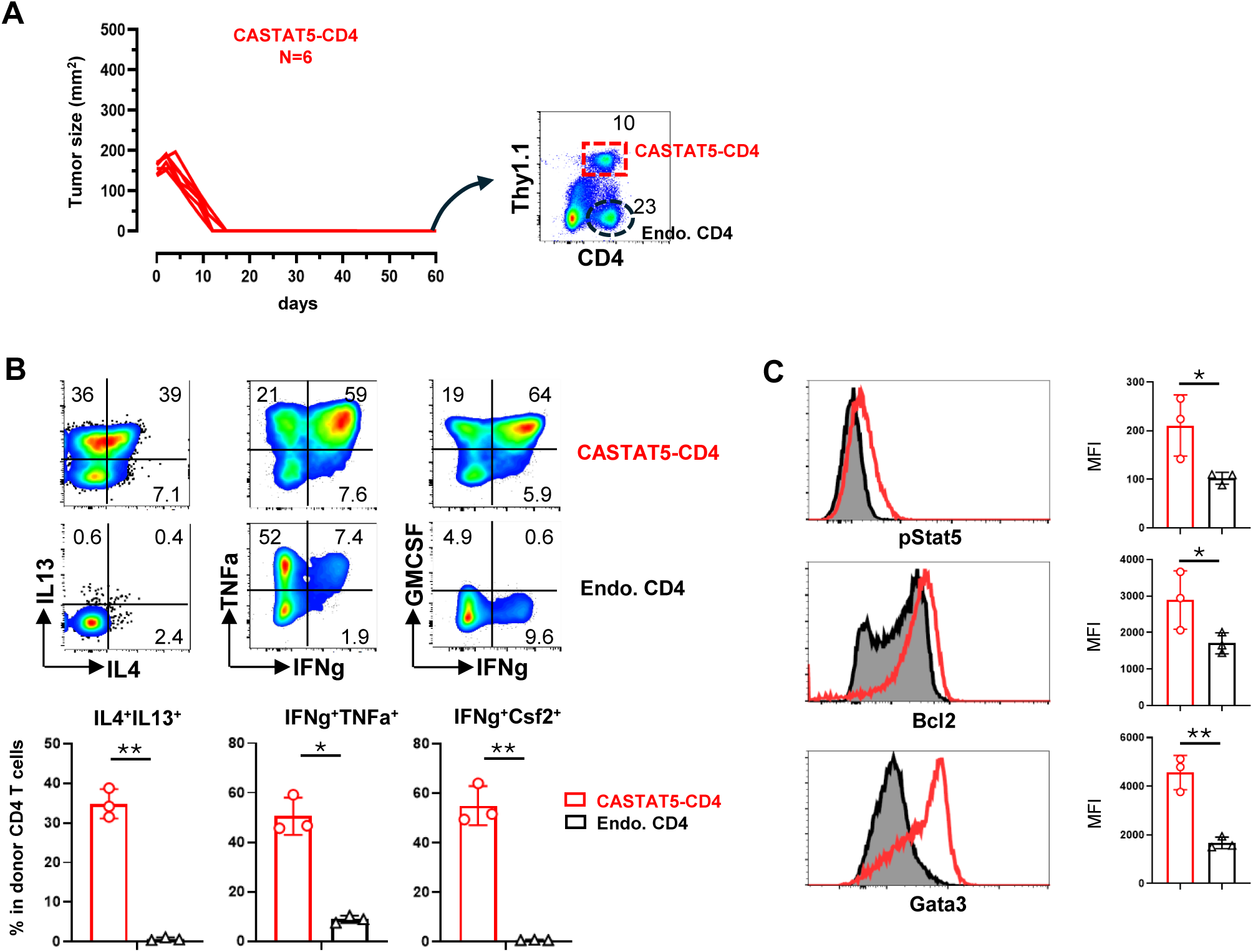
CASTAT5-CD4 T cells retain polyfunctionality long after tumor eradication. A. C5-CD4 donor T cells remain readily detectable in recipient mice after tumor eradication. The experimental setup is described in Figure 1. Spleens were collected from mice that remained tumor-free for more than 60 days after infusion of C5-CD4 T cells. A representative dot plot shows the detection of donor CD4^+^ T cells. B. Polyfunctionality of donor CD4^+^ T cells assessed by ICS, as reflected by the co-expression of multiple cytokines. Representative dot plots are shown, with numbers indicating the percentage of cells in the corresponding quadrant. Endogenous CD4^+^ T cells were included for comparison and to establish the gating strategy. Quantification is summarized in the corresponding bar graphs as mean ± SE (n = 3 mice per group). C. Expression of pStat5, Bcl2, and Gata3 in persistent C5-CD4 T cells assessed by intracellular staining. Representative histograms show overlay profiles of the indicated molecules in C5-CD4 T cells and endogenous CD4^+^ T cells. Mean fluorescence intensities (MFIs) are summarized in the corresponding bar graphs as mean ± SE (n = 3 mice per group). *p < 0.05, **p < 0.01.

**Supplemental Figure 2.**
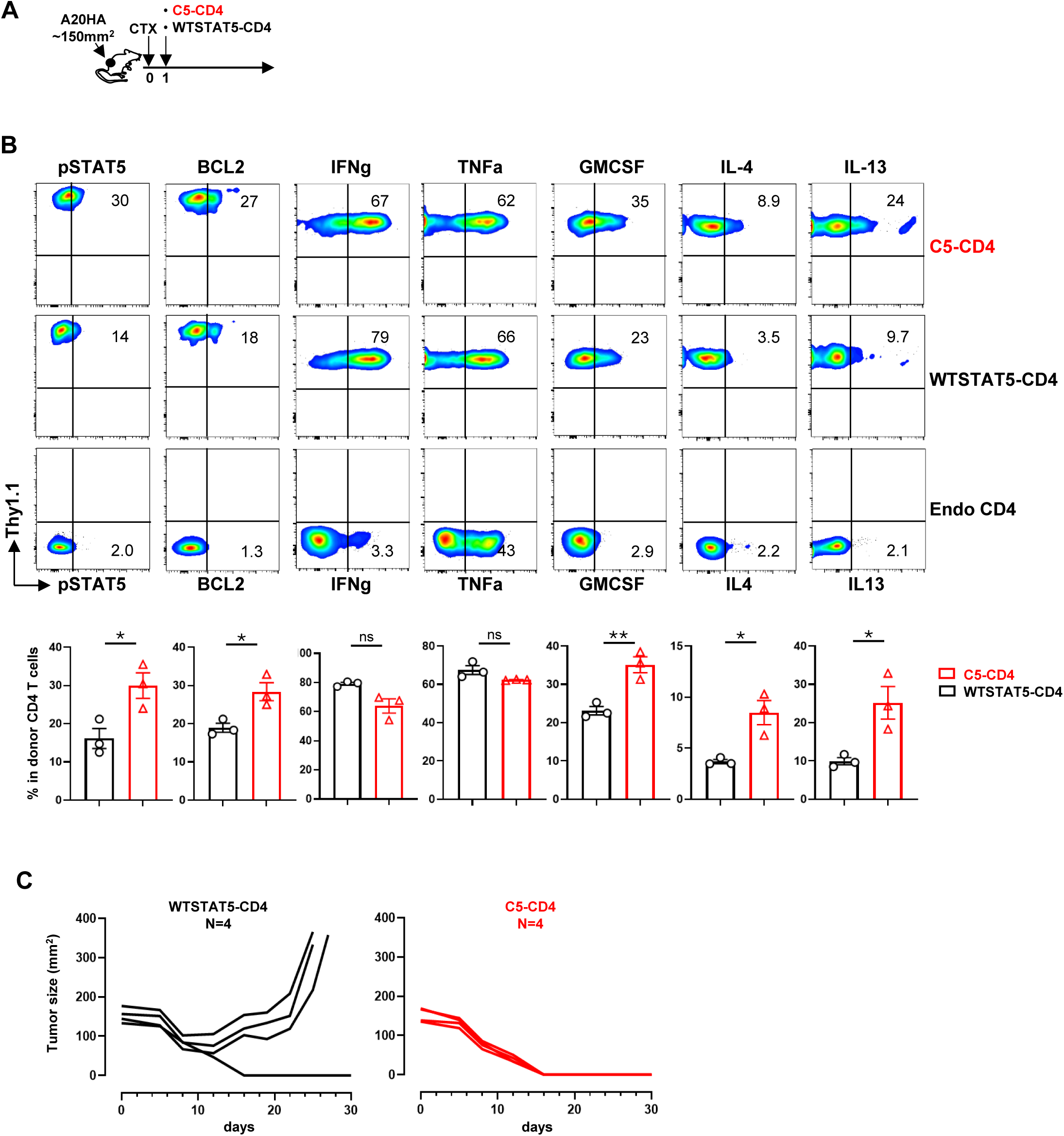
Overexpression of wild-type Stat5 does not recapitulate the CASTAT5 phenotype. **A.** Schematic of the experimental design. **B.** pStat5, Bcl2 and cytokine profile of donor CD4^+^ T cells assessed by ICS. Following adoptive transfer of C5-CD4 and WTCASTAT5-CD4 T cells, peripheral blood was collected by tail vein 7 days later for ICS. Representative dot plots are shown for each condition, with numbers indicating the percentages of cytokine-producing cells among donor CD4^+^ T cells. Endogenous CD4^+^ T cells were included to establish the gating strategy. Quantification is summarized in the corresponding bar graphs as mean ± SE (n = 3 mice per group). **C.** Tumor growth curves for individual mice in each treatment group. ns, not significant; *, p < 0.05; **, p < 0.01.

**Supplemental Figure 3.**
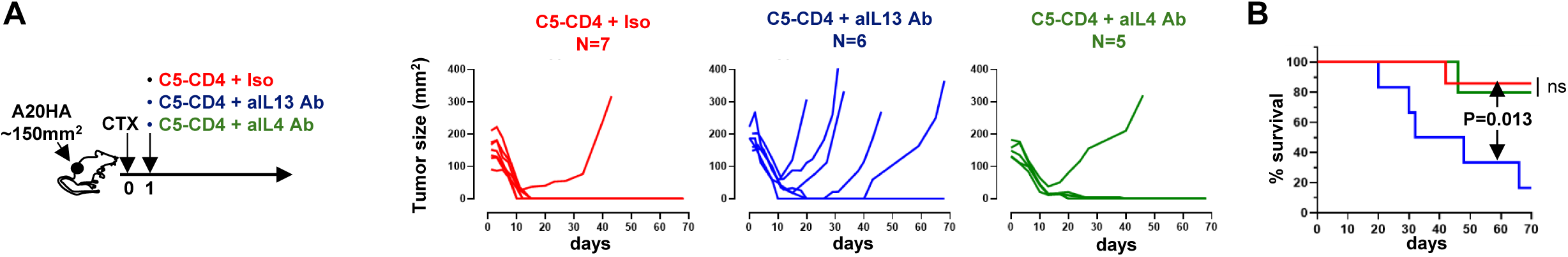
IL13 but not IL4 neutralization diminishes the antitumor efficacy of C5-CD4 T cells. **A.** A20HA-bearing mice received adoptive transfer of C5-CD4 T cells and were treated with an isotype control, an anti-IL13, or an anti-IL4 neutralizing antibody, as indicated in the schematic. Tumor growth curves for individual mice are shown. **B.** Kaplan-Meier survival analysis of the mice shown in A.

**Supplemental Figure 4.**
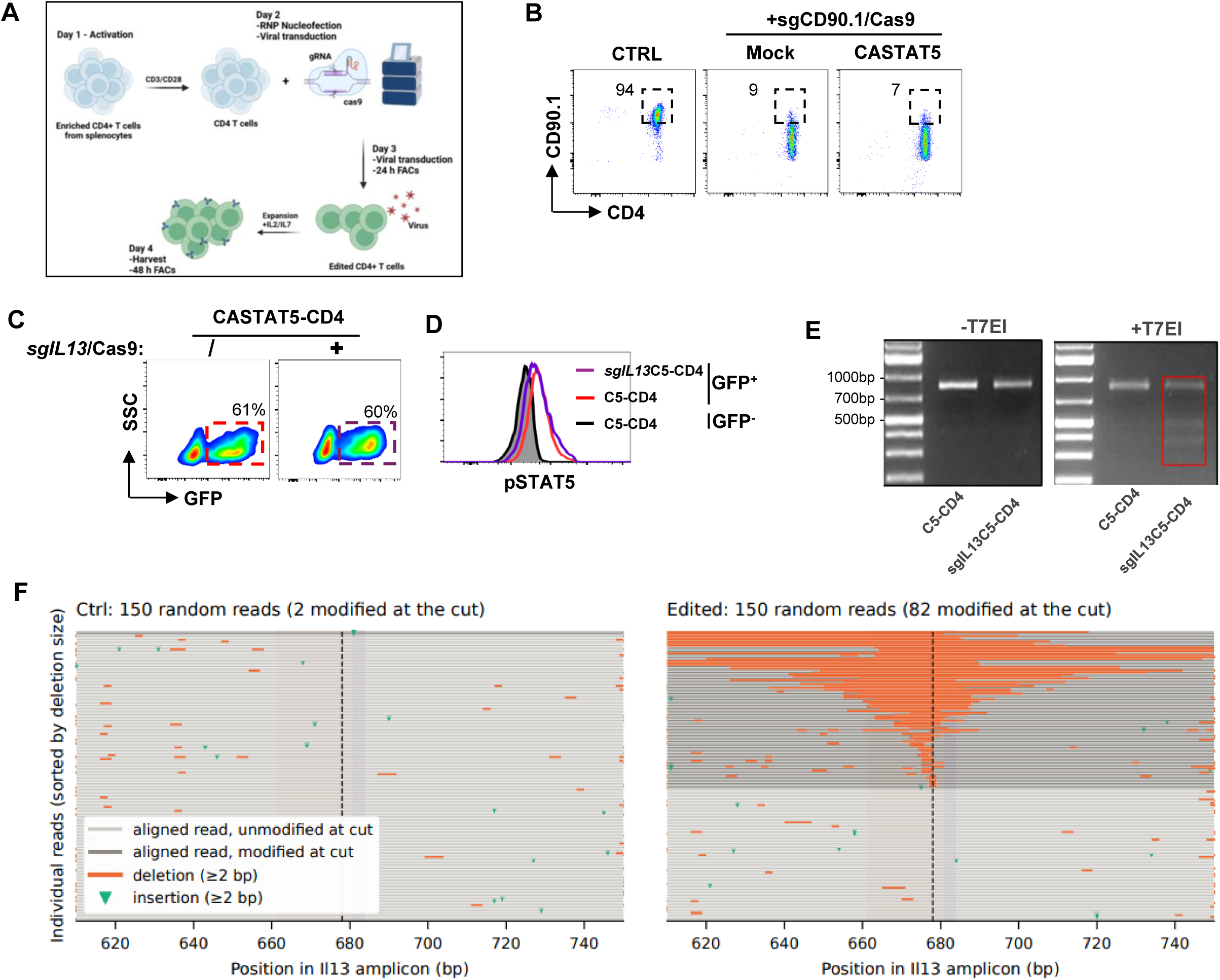
Validation of the CRISPR/RNP approach for targeted gene editing in T cells. A. Schematic of the experimental procedures. Briefly, CD4+ T cells isolated from splenocytes of 6.5 TCR-Tg mice on a Thy1.1 (CD90.1) background were stimulated overnight with T-cell activation Dynabeads. The following day, T cells were nucleofected with preformed RNP complexes containing Cas9 and gene-specific sgRNA, followed by viral transduction with CASTAT5. T cells were transduced again with CASTAT5-carrying virus the next day and expanded for an additional day before being harvested for analysis or infused into recipient mice. B. Assessment of gene-editing efficiency using the CRISPR/RNP approach. Activated CD4+ T cells were either left unmanipulated (CTRL) or nucleofected with CD90.1-targeting RNP, followed by mock transduction or CASTAT5 viral transduction. Dot plots show CD90.1 expression in CD4+ T cells under the indicated conditions, with numbers indicating the percentages of CD90.1+ cells among CD4+ T cells. C. IL13-targeted gene editing by RNP does not affect CASTAT5 transduction efficiency in CD4+ T cells. Activated CD4+ T cells were either left unnucleofected or nucleofected with IL13-targeting RNP, followed by CASTAT5 viral transduction. Representative dot plots show GFP expression in CD4+ T cells under the indicated conditions. D. pSTAT5 levels in the cells shown in (C) assessed by intracellular staining. E. Validation of targeted gene editing by T7 endonuclease I (T7E1) digestion assay. GFP+ cells shown in (C) were enriched by cell sorting. Genomic DNA was extracted from the sorted cells and used as the template for PCR amplification of the IL13 genomic region containing the IL13.sgRNA target sequence. PCR products were either left undigested or digested with T7E1 for 2 h before electrophoresis. The gels show undigested (left) and T7E1-digested (right) PCR products. The appearance of additional shorter bands in the sgIl13C5-CD4 sample (red box) indicates the occurrence of targeted gene editing. F. Confirmation of IL13-targeted gene editing by Nanopore sequencing. PCR products covering the targeted region were subjected to Nanopore sequencing. Representative plots demonstrate deletions at the targeted *Il13* gene locus.

**Supplemental Figure 5.**
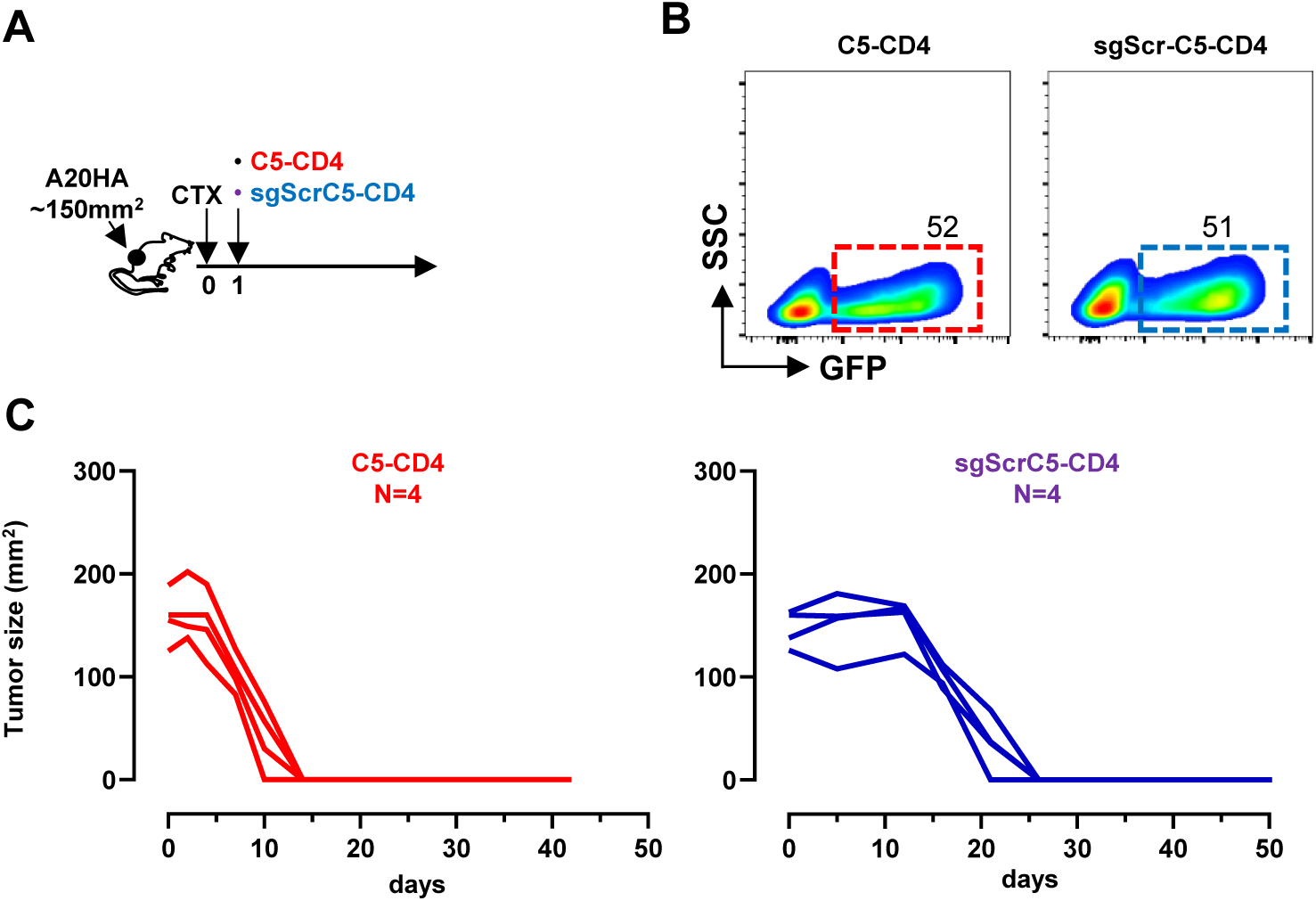
The RNP nucleofection procedure per se does not alter the antitumor activity of C5-CD4 T cells. **A.** Schematic of the experimental design. **B.** CASTAT5 transduction efficiency assessed by GFP expression. Activated HA-specific CD4^+^ T cells were either left unnucleofected or nucleofected with scramble sgRNA, followed by CASTAT5 viral transduction. Representative dot plots show GFP expression in CD4^+^ T cells under the indicated conditions, with numbers indicating the percentages of GFP^+^ cells. GFP^+^ cells were isolated by flow sorting and used for adoptive transfer. **C.** Tumor growth curves for individual mice in each treatment group.

**Supplemental Figure 6.**
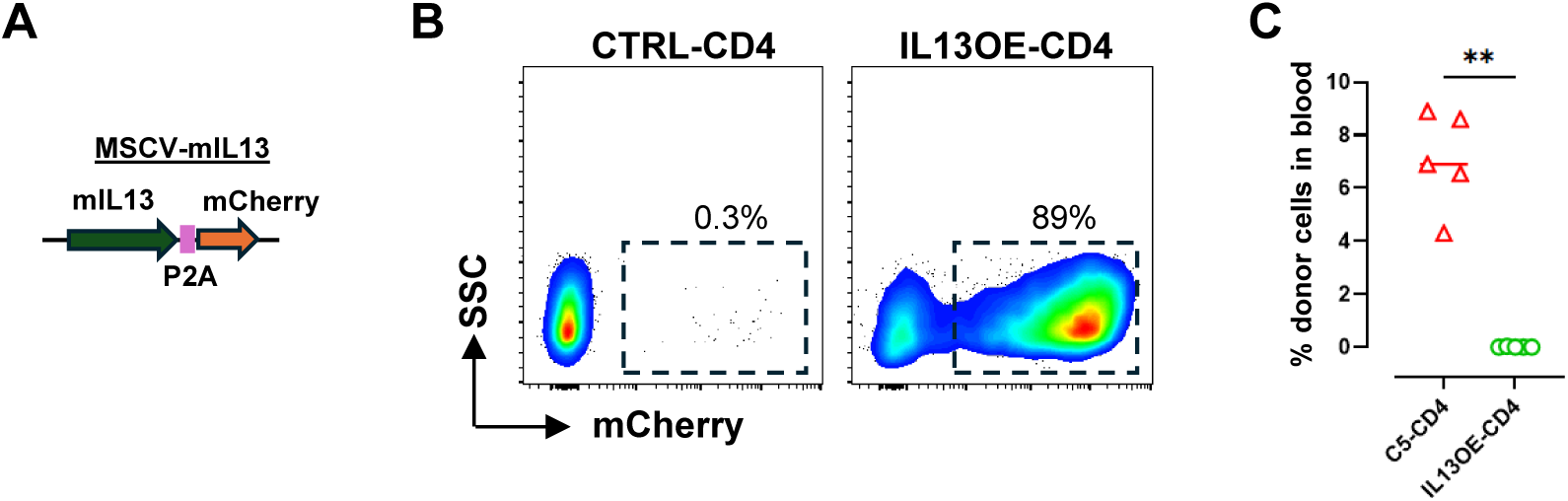
Generation of IL13OE-CD4 T cells. **A.** Schematic of the retroviral vector encoding murine IL13 followed by a P2A-linked mCherry reporter. **B.** Activated HA-specific CD4^+^ T cells were mock-transduced or transduced with viruses carrying IL13-mCherry. Transduction efficiency was measured by percent of mCherry-positive cells. Transduced cells were used for adoptive transfer experiments depicted in Fig. 3H. **C.** Frequencies of donor CD4^+^ T cells in peripheral blood 21 days after adoptive transfer. **, p < 0.01

**Supplemental Figure 7.**
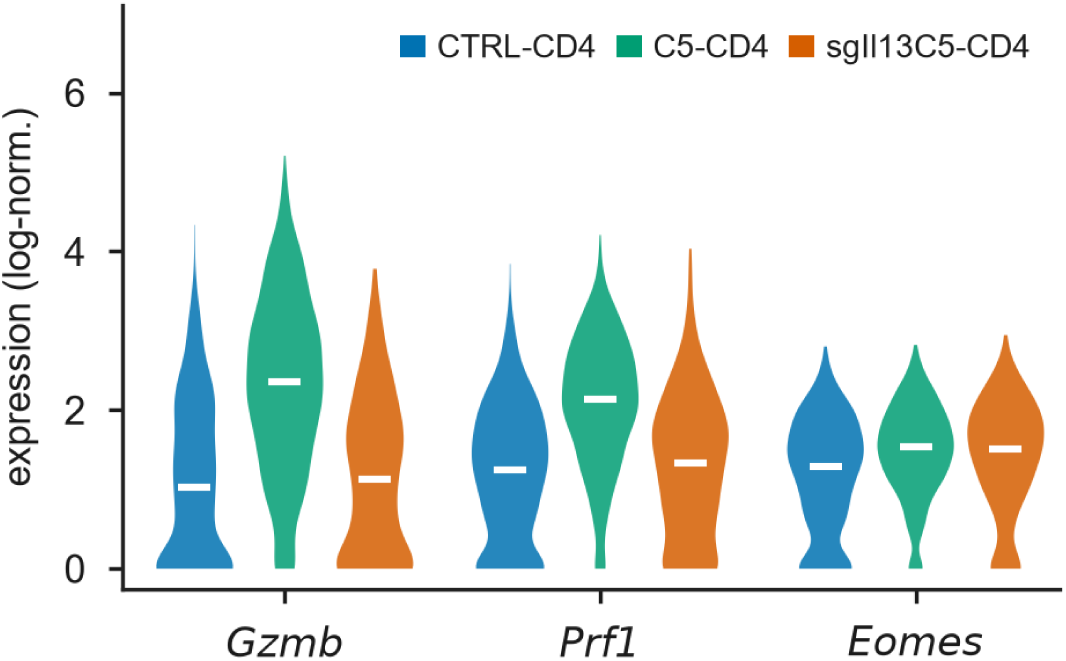
IL13 deficiency is associated with phenotypical changes in NK cells. Expression of cytotoxic effector genes (*Gzmb*, *Prf1* and *Eomes*) in tumor-infiltrating NK cells. Violin plots show log-normalized gene expression, with white bars indicating medians.

**Supplemental Figure 8.**
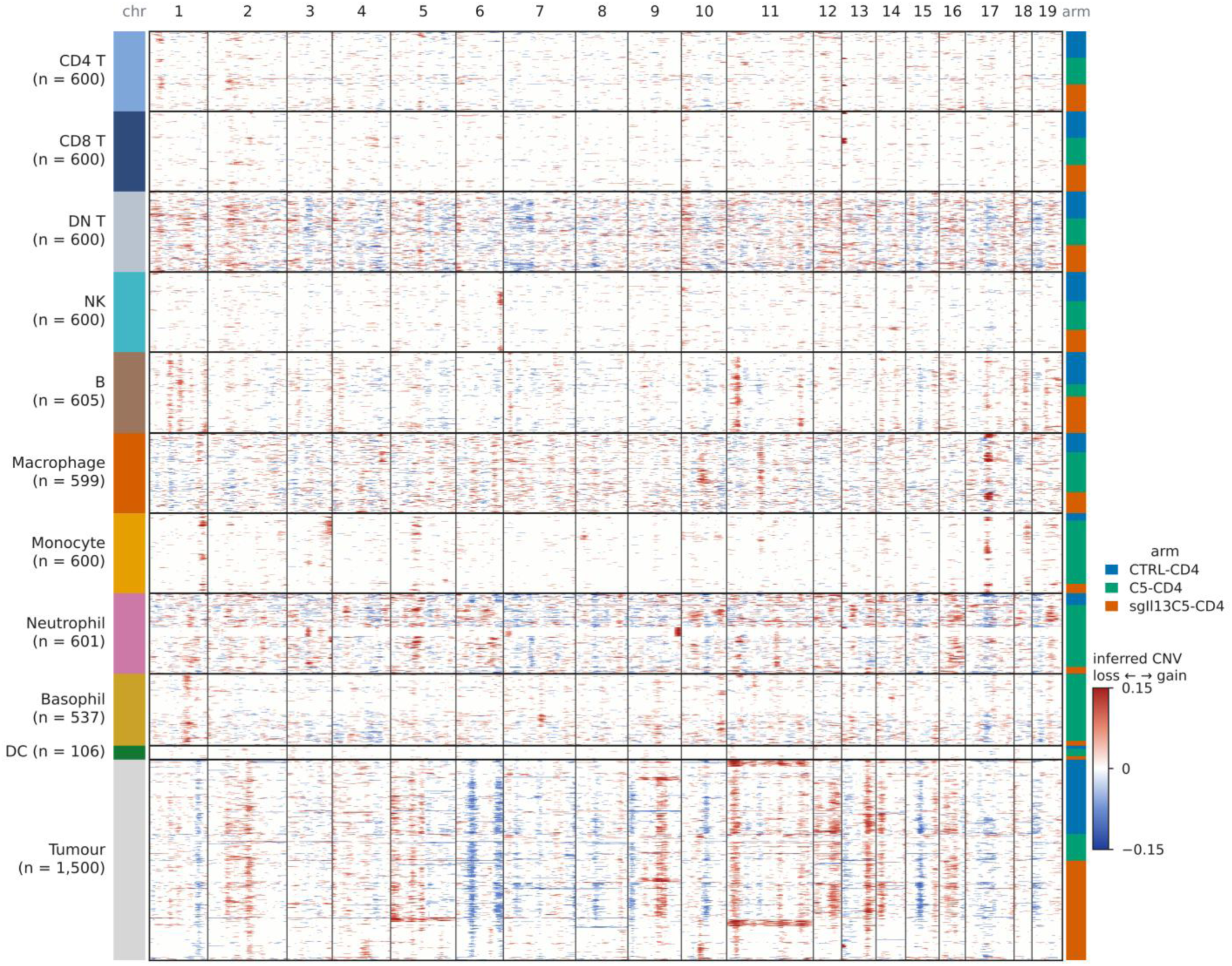
**Inferred copy-number alterations distinguish A20 lymphoma cells from host immune populations**. The InferCNV heatmap shows inferred copy-number variation (CNV) profiles of tumor and immune cells from CTRL-CD4, C5-CD4, and sgIl13C5-CD4 tumors. A total of 12,805 cells were included, comprising all annotated tumor and B cells and up to 500 cells per treatment group from each remaining cell type. Rows represent individual cells grouped by cell-type annotation (left color bar; colors as in Fig. 4C), treatment group (right color bar), and hierarchical clustering. Columns represent 100-gene windows ordered along the mm10 autosomes. Colors indicate smoothed expression relative to host immune reference populations (blue, inferred loss; red, inferred gain; values clipped at ±0.15). For visualization, cell populations were downsampled to a maximum of 600 cells per group, except tumor cells (1,500 cells). Annotated tumor cells displayed a shared CNV pattern across treatment groups, characterized by inferred losses on chromosomes 6, 9, 10, and 15 and gains on chromosomes 9, 12, 13, and 14, which was absent from host immune populations, including B cells.

**Supplemental Figure 9.**
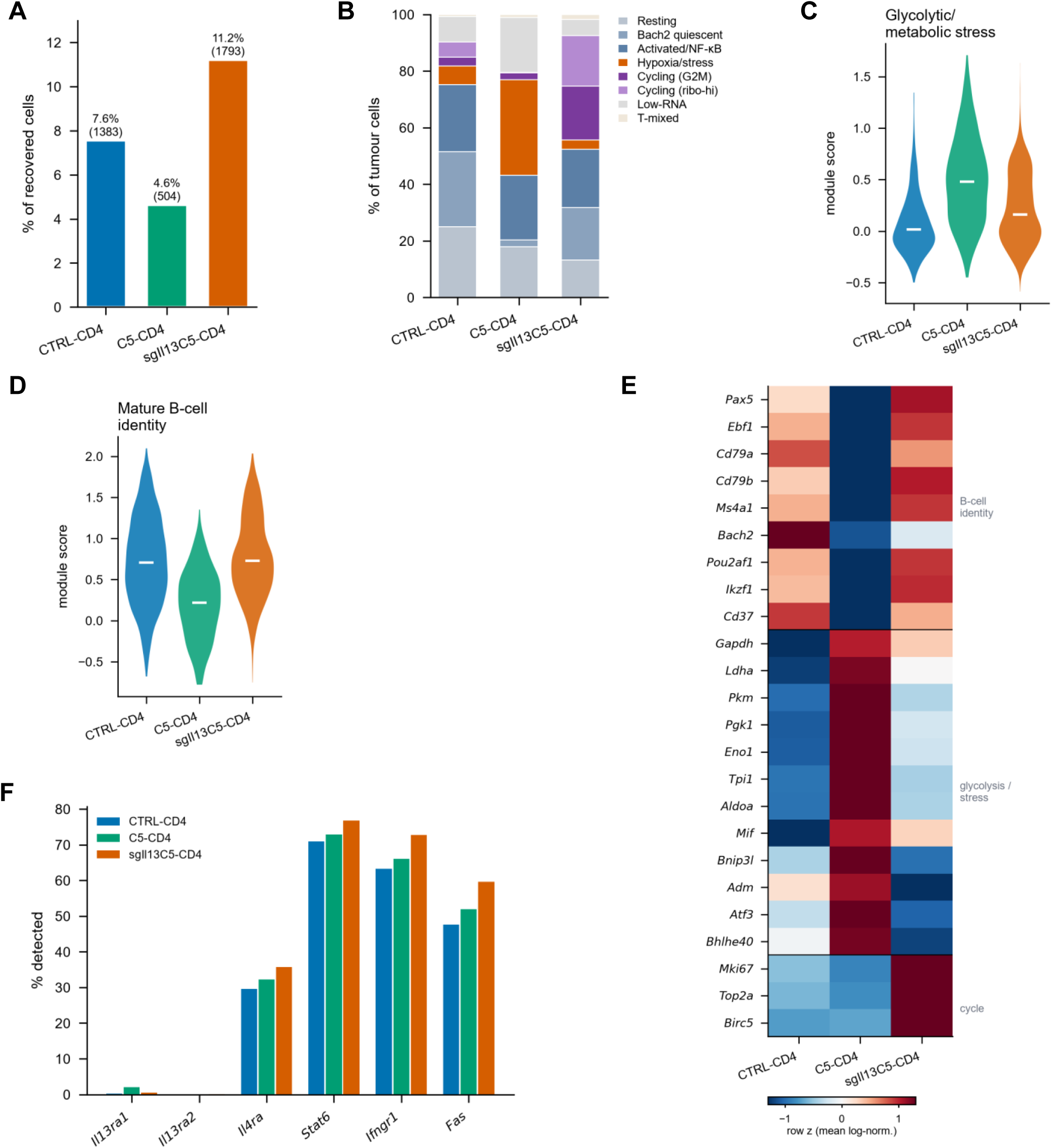
IL13 deficiency in C5-CD4 T cells attenuates transcriptional remodeling of A20 tumor cells. scRNA-seq analysis of tumor samples are described in Figure 4. A20HA tumor cells were identified by level-3 cell-type annotation and independently re-clustered. **A.** Relative abundance of tumor cells among all recovered cells in each condition. Numbers above bars indicate recovered tumor-cell counts. Percentages represent capture fractions rather than absolute tumor burden. **B.** Distribution of tumor cells across eight transcriptional states identified by Leiden clustering (resolution 0.5) using 2,000 highly variable genes, 20 principal components and 20 nearest neighbors. Immunoglobulin, ribosomal and mitochondrial genes were excluded from clustering. States were annotated on the basis of marker-gene expression as resting, *Bach2*-high quiescent, activated/NF-κB, hypoxia/stress, cycling (G2/M), cycling/ribosomal-high, low- RNA and T-cell-mixed. **C.** Glycolytic/metabolic-stress module scores based on *Gapdh*, *Tpi1*, *Pkm*, *Ldha*, *Pgk1*, *Eno1*, *Pgam1*, *Aldoa*, *Gpi1*, *Pfkp*, *Pfkl*, *Hk2* and *Mif*. Mean scores were 0.08, 0.50 and 0.22 in CTRL-CD4, C5-CD4 and sgIl13C5-CD4 tumors, respectively. **D.** Mature B-cell identity module scores based on *Pax5*, *Ebf1*, *Cd79a*, *Cd79b*, *Ms4a1*, *Cd37*, *Bach2*, *Pou2af1*, *Ikzf1*, *Cd19* and *Bank1*. Corresponding mean scores were 0.72, 0.19 and 0.77. In C and D, violin plots show single-cell score distributions, with white bars indicating medians. Module scores were calculated from log-normalized expression relative to matched control genes using Scanpy score_genes. **E.** Heatmap showing mean log-normalized expression of genes associated with B-cell identity, glycolysis/metabolic stress and cell-cycle progression across the three conditions. Expression values were standardized for each gene (row z-score). **F.** Percentage of tumor cells with detectable expression of the cytokine-receptor and death-receptor genes *Il13ra1*, *Il13ra2*, *Il4ra*, *Stat6*, *Ifngr1* and *Fas*. *Il13ra1* and *Il13ra2* transcripts were detected in ≤2% of tumor cells across all conditions, indicating limited detectable expression of these receptor subunits. Expression values were normalized to 10,000 counts per cell and log-transformed [log1p(counts/library size × 10⁴)]. As each condition was represented by a single pooled sample, all comparisons are descriptive and no statistical inference across biological replicates was performed.

**Supplemental Figure 10.**
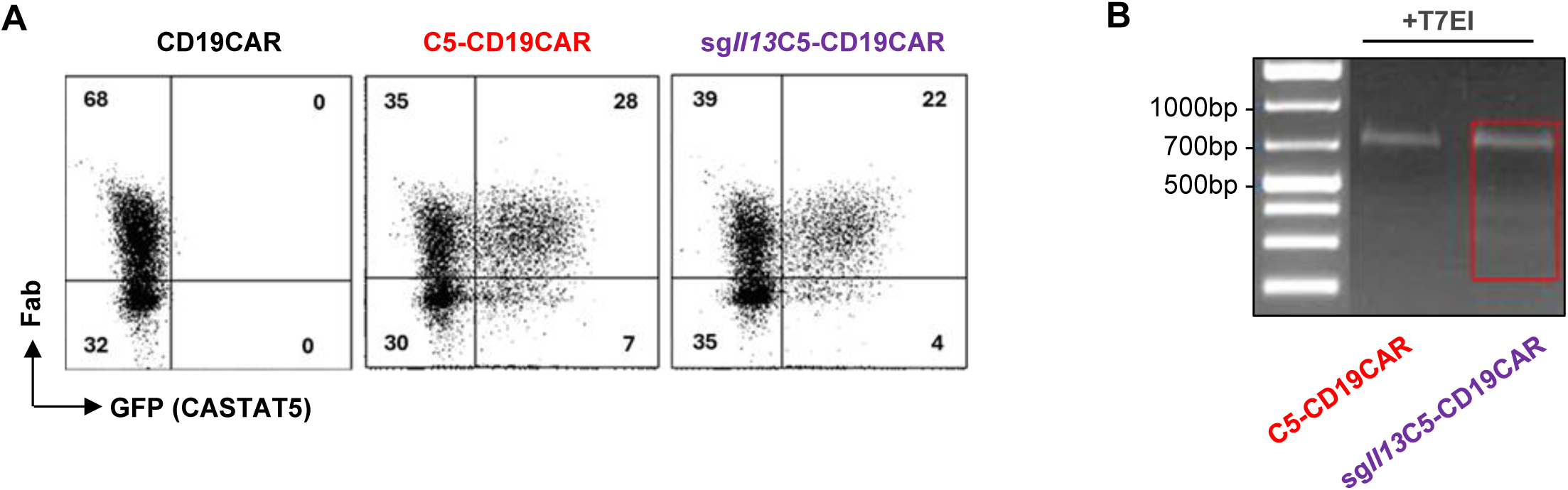
Validation of IL13-targeted gene editing in C5-CD19CAR-T cells. **A.** Viral transduction efficiency in T cells. Total T cells were purified from CD45.1 congenic mice and transduced to express CD19CAR or co-express CD19CAR and CASTAT5. A subset of CASTAT5/CD19CAR-cotransduced T cells was nucleofected with sg*Il13* RNP. Representative dot plots show CD19CAR and CASTAT5 expression in transduced T cells. CD19CAR was detected by anti-Fab staining, and CASTAT5 expression was assessed by GFP expression. Numbers indicate the percentages of cells in the corresponding quadrants. **B.** Assessment of targeted gene editing by T7E1 digestion assay. Genomic DNA was extracted from the indicated T-cell populations and subjected to PCR amplification as described in Supplemental Figure 4E. The gel shows the results of T7E1 digestion of the PCR products.

## References

1. E. Tran et al., Cancer immunotherapy based on mutation-specific CD4+ T cells in a patient with epithelial cancer. Science 344, 641–645 (2014).

2. I. Liadi et al., Individual Motile CD4(+) T Cells Can Participate in Efficient Multikilling through Conjugation to Multiple Tumor Cells. Cancer Immunol Res 3, 473–482 (2015).

3. D. Wang, et al., Glioblastoma-targeted CD4+ CAR T cells mediate superior antitumor activity. JCI Insight 3, (2018).

4. S. Agarwal et al., In Vivo Generation of CAR T Cells Selectively in Human CD4(+) Lymphocytes. Mol Ther 28, 1783–1794 (2020).

5. K. Hung et al., The central role of CD4(+) T cells in the antitumor immune response. J Exp Med 188, 2357–2368 (1998).

6. N. N. Hunder et al., Treatment of metastatic melanoma with autologous CD4+ T cells against NY-ESO-1. N Engl J Med 358, 2698–2703 (2008).

7. A. Corthay et al., Primary antitumor immune response mediated by CD4+ T cells. Immunity 22, 371–383 (2005).

8. A. Perez-Diez et al., CD4 cells can be more efficient at tumor rejection than CD8 cells. Blood 109, 5346–5354 (2007).

9. M. Liu et al., TGF-beta suppresses type 2 immunity to cancer. Nature 587, 115–120 (2020).

10. B. Kruse et al., CD4(+) T cell-induced inflammatory cell death controls immune-evasive tumours. Nature 618, 1033–1040 (2023).

11. G. Espinosa-Carrasco et al., Intratumoral immune triads are required for immunotherapy-mediated elimination of solid tumors. Cancer Cell 42, 1202–1216 e1208 (2024).

12. Q. Lian et al., CD4(+) T cells impair tumor growth through IL-3 and TNF-dependent vascular damage. Science 392, eads7910 (2026).

13. F. Zhang Perkins et al., Synthetic long peptide and DNA personalized cancer vaccines induce robust neoantigen-specific T cell responses in pancreatic cancer. Sci Adv 12, eaei1190 (2026).

14. C. J. Maine et al., Self-Replicating RNAs Drive Protective Anti-tumor T Cell Responses to Neoantigen Vaccine Targets in a Combinatorial Approach. Mol Ther 29, 1186–1198 (2021).

15. N. Markosyan, R. H. Vonderheide, Developing neoantigen cancer vaccines: where are we now? Nat Commun 17, (2026).

16. J. L. Tanyi et al., Personalized cancer vaccine strategy elicits polyfunctional T cells and demonstrates clinical benefits in ovarian cancer. NPJ Vaccines 6, 36 (2021).

17. S. A. Quezada et al., Tumor-reactive CD4(+) T cells develop cytotoxic activity and eradicate large established melanoma after transfer into lymphopenic hosts. J Exp Med 207, 637–650 (2010).

18. Y. Xie et al., Naive tumor-specific CD4(+) T cells differentiated in vivo eradicate established melanoma. J Exp Med 207, 651–667 (2010).

19. Z. C. Ding et al., Polyfunctional CD4(+) T cells are essential for eradicating advanced B-cell lymphoma after chemotherapy. Blood 120, 2229–2239 (2012).

20. J. Rossi et al., Preinfusion polyfunctional anti-CD19 chimeric antigen receptor T cells are associated with clinical outcomes in NHL. Blood 132, 804–814 (2018).

21. A. L. Huff, et al., CD4 T cell-activating neoantigens enhance personalized cancer vaccine efficacy. JCI Insight 8, (2023).

22. Z. C. Ding, et al., Persistent STAT5 activation reprograms the epigenetic landscape in CD4(+) T cells to drive polyfunctionality and antitumor immunity. Sci Immunol 5, (2020).

23. D. G. DeNardo, P. Andreu, L. M. Coussens, Interactions between lymphocytes and myeloid cells regulate pro- versus anti-tumor immunity. Cancer Metastasis Rev 29, 309– 316 (2010).

24. S. L. Shiao, A. P. Ganesan, H. S. Rugo, L. M. Coussens, Immune microenvironments in solid tumors: new targets for therapy. Genes Dev 25, 2559–2572 (2011).

25. E. Jou, Type 1 and type 2 cytokine-mediated immune orchestration in the tumour microenvironment and their therapeutic potential. Explor Target Antitumor Ther 4, 474– 497 (2023).

26. M. Terabe et al., NKT cell-mediated repression of tumor immunosurveillance by IL-13 and the IL-4R-STAT6 pathway. Nat Immunol 1, 515–520 (2000).

27. C. Aspord et al., Breast cancer instructs dendritic cells to prime interleukin 13-secreting CD4+ T cells that facilitate tumor development. J Exp Med 204, 1037–1047 (2007).

28. D. G. DeNardo et al., CD4(+) T cells regulate pulmonary metastasis of mammary carcinomas by enhancing protumor properties of macrophages. Cancer Cell 16, 91–102 (2009).

29. M. A. Hallett, K. T. Venmar, B. Fingleton, Cytokine stimulation of epithelial cancer cells: the similar and divergent functions of IL-4 and IL-13. Cancer Res 72, 6338–6343 (2012).

30. R. I. Tepper, R. L. Coffman, P. Leder, An eosinophil-dependent mechanism for the antitumor effect of interleukin-4. Science 257, 548–551 (1992).

31. O. S. Blomberg et al., IL-5-producing CD4(+) T cells and eosinophils cooperate to enhance response to immune checkpoint blockade in breast cancer. Cancer Cell 41, 106–123 e110 (2023).

32. T. Nishimura et al., Distinct role of antigen-specific T helper type 1 (Th1) and Th2 cells in tumor eradication in vivo. J Exp Med 190, 617–627 (1999).

33. K. B. Lorvik et al., Adoptive Transfer of Tumor-Specific Th2 Cells Eradicates Tumors by Triggering an In Situ Inflammatory Immune Response. Cancer Res 76, 6864–6876 (2016).

34. D. Jacenik, I. Karagiannidis, E. J. Beswick, Th2 cells inhibit growth of colon and pancreas cancers by promoting anti-tumorigenic responses from macrophages and eosinophils. Br J Cancer 128, 387–397 (2023).

35. Z. Bai et al., Single-cell CAR T atlas reveals type 2 function in 8-year leukaemia remission. Nature 634, 702–711 (2024).

36. M. Yamashita et al., Identification of a conserved GATA3 response element upstream proximal from the interleukin-13 gene locus. J Biol Chem 277, 42399–42408 (2002).

37. P. E. Fields, G. R. Lee, S. T. Kim, V. V. Bartsevich, R. A. Flavell, Th2-specific chromatin remodeling and enhancer activity in the Th2 cytokine locus control region. Immunity 21, 865–876 (2004).

38. H. Nagashima et al., Remodeling of Il4-Il13-Il5 locus underlies selective gene expression. Nat Immunol 25, 2220–2233 (2024).

39. E. I. Frolova et al., Linkage mapping of the human CSF2 and IL3 genes. Proc Natl Acad Sci U S A 88, 4821–4824 (1991).

40. A. V. Villarino, et al., A central role for STAT5 in the transcriptional programing of T helper cell metabolism. Sci Immunol 7, eabl9467 (2022).

41. A. Seki, S. Rutz, Optimized RNP transfection for highly efficient CRISPR/Cas9-mediated gene knockout in primary T cells. J Exp Med 215, 985–997 (2018).

42. M. Terabe, J. M. Park, J. A. Berzofsky, Role of IL-13 in regulation of anti-tumor immunity and tumor growth. Cancer Immunol Immunother 53, 79–85 (2004).

43. B. Feng et al., The type 2 cytokine Fc-IL-4 revitalizes exhausted CD8(+) T cells against cancer. Nature 634, 712–720 (2024).

44. J. J. Melenhorst et al., Decade-long leukaemia remissions with persistence of CD4(+) CAR T cells. Nature 602, 503–509 (2022).

45. I. Xhangolli et al., Single-cell Analysis of CAR-T Cell Activation Reveals A Mixed T(H)1/T(H)2 Response Independent of Differentiation. Genomics Proteomics Bioinformatics 17, 129–139 (2019).

46. J. Kirberg et al., Thymic selection of CD8+ single positive cells with a class II major histocompatibility complex-restricted receptor. J Exp Med 180, 25–34 (1994).

47. J. N. Kochenderfer, Z. Yu, D. Frasheri, N. P. Restifo, S. A. Rosenberg, Adoptive transfer of syngeneic T cells transduced with a chimeric antigen receptor that recognizes murine CD19 can eradicate lymphoma and normal B cells. Blood 116, 3875–3886 (2010).

48. C. S. Ross-Innes et al., Differential oestrogen receptor binding is associated with clinical outcome in breast cancer. Nature 481, 389–393 (2012).

49. G. Yu, L. G. Wang, Q. Y. He, ChIPseeker: an R/Bioconductor package for ChIP peak annotation, comparison and visualization. Bioinformatics 31, 2382–2383 (2015).

50. N. C. Durand et al., Juicer Provides a One-Click System for Analyzing Loop-Resolution Hi-C Experiments. Cell Syst 3, 95–98 (2016).

51. Y. Hao et al., Dictionary learning for integrative, multimodal and scalable single-cell analysis. Nat Biotechnol 42, 293–304 (2024).

52. A. P. Patel et al., Single-cell RNA-seq highlights intratumoral heterogeneity in primary glioblastoma. Science 344, 1396–1401 (2014).

53. S. Jin et al., Inference and analysis of cell-cell communication using CellChat. Nat Commun 12, 1088 (2021).

54. B. Van de Sande et al., A scalable SCENIC workflow for single-cell gene regulatory network analysis. Nat Protoc 15, 2247–2276 (2020).

