## Supplemental Table I for "IL13 critically contributes to the antitumor activity of polyfunctional CD4^+^ T cells driven by constitutively active Stat5"

**Table I. Reagents used for *Il13* sgRNA/Cas9 RNP-mediated gene editing and validation.**

| Reagent name | Sequences |
|---|---|
| *Il13* crRNA1 | rGrArCrCrCrUrUrArArGrGrArGrCrUrUrArUrUrGrGrUrUrUrUrArGrArGrCrUrArUrGrCrU |
| *Il13* crRNA2 | rGrArGrGrCrCrArUrGrCrArArUrArUrCrCrUrCrUrGrUrUrUrUrArGrArGrCrUrArUrGrCrU |
| Alt-R™ CRISPR-Cas9 tracrRNA | |
| Alt-R® CRISPR-Cas9 Negative Control crRNA #1 | |
| Alt-R® CRISPR-Cas9 Negative Control crRNA #2 | |
| *Il13* gRNA-1 PCR primer-F | TTCTACAGCTCCCTGGTTCT |
| *Il13* gRNA-1 PCR primer-R | GAAGAGTCCCGAGCTAGTAGAA |
| *Il13* gRNA-2 PCR primer-F | CCTAATGCTGTGACCCTTAACT |
| *Il13* gRNA-2 PCR primer-R | GGGATGGTCTCTCCTCATTAGA |
